# Phenotype prediction using biologically interpretable neural networks on multi-cohort multi-omics data

**DOI:** 10.1101/2023.04.16.537073

**Authors:** Arno van Hilten, Jeroen van Rooij, BIOS consortium, M. Arfan Ikram, Wiro. J. Niessen, Joyce. B.J. van Meurs, Gennady V. Roshchupkin

## Abstract

Integrating multi-omics data into predictive models has the potential to enhance accuracy, which is essential for precision medicine. In this study, we developed interpretable predictive models for multi-omics data by employing neural networks informed by prior biological knowledge, referred to as visible networks. These neural networks offer insights into the decision-making process and can unveil novel perspectives on the underlying biological mechanisms associated with traits and complex diseases. We tested the performance, interpretability, and generalizability for inferring smoking status, subject age and LDL levels using genome-wide RNA-expression and CpG methylation data from blood of the BIOS consortium(4 population cohorts, N_total=2940). In a cohort-wise cross validation setting, the consistency of the diagnostic performance and interpretation was assessed.

Performance was consistently high for predicting smoking status with an overall mean AUC of 0.95 (95% CI, 0.90 - 1.00) and interpretation revealed the involvement of well-replicated genes such as *AHRR, GPR15* and *LRRN3*. LDL-level predictions only generalized in a single cohort with an R^2^ of 0.07 (95% CI, 0.05 - 0.08). Age was infered with a mean error of 5.16 (95% CI, 3.97 - 6.35) years with the genes *COL11A2, AFAP1, OTUD7A, PTPRN2, ADARB2* and *CD34* consistently predictive. In general, we found that using multi-omics networks improved performance, stability and generalizability compared to interpretable single omic networks.

We believe that visible neural networks have great potential for multi-omics analysis; they combine multi-omic data elegantly, are interpretable, and generalize well to data from different cohorts.

## Introduction

Over the last decades, association studies have uncovered numerous genes and CpGs to be associated with hundreds of traits and diseases^1^. This has led to tools for identifying high risk individuals and biomarkers for early disease detection. For example, blood-based methylation biomarkers are currently used for early diagnosis for various forms of cancer^2,3^. However, for most complex diseases and traits, the combined effects, within and between different omics types, is still largely unexplored. For a more comprehensive understanding of human health and diseases and for more accurate prediction models, it is therefore necessary to study omic types in relation to one another. Thanks to recent technological improvements for high throughput sequencing and arrays technologies, the acquisition of multi-omics datasets has become more feasible, providing opportunities for new multi-omics analysis tools^4,5^.

Recently, novel statistical frameworks and machine learning techniques have been published that integrate multi-omics data in a single analysis^6,7^. These studies show the potential for multi-omics analysis to improve prediction for various disorders while providing insight into the disease biology^4,8^. Integrating different types of omics data in a single analysis is a challenging task, as each type has different, procedures, preprocessing steps and analytical requirements^9^. Combining omics data presents additional challenges, as each omic has unique dimensions, and it is essential to consider correlation structures both within and between the different omics types. Thus, for the combined analysis of multiple omics types, methods need to be flexible and be able to deal with the high dimensionality of these datasets.

Neural networks have demonstrated such flexibility and have been widely successful in fields such as image classification^10^, speech recognition^11^, and protein modelling^12^. In contrast to most tasks in image analysis and speech recognition, the focus of multi-omics frameworks is not only on predictive performance but also in understanding the underlying etiology. To facilitate this, a new field in machine learning, coined visible machine learning^13^ emerged, in which prior biological knowledge is embedded in a neural network’s architecture to create interpretable neural networks^14–17^. Recent examples of these kinds of neural networks applied in genomics are GenNet^18^ and P-net^19^. In the GenNet framework, gene and pathway annotations were used to create interpretable neural networks for genetic risk prediction from genotype. In P-net, methylation, gene expression and copy number variants were fed to an interpretable neural network to differentiate between primary or metastatic prostate cancers. Other examples include, PasNet^20^, which integrated pathways information to predict survival for glioblastoma multiforme, a primary brain cancer. DrugCell^21^ integrated Gene Ontology knowledge in a network to predict drug response for various cancers and ParsVNN^22^ continued on this work and pruned the network for increased performance and better interpretability.

In this study, we create visible neural networks to analyze multi-omics data in a single analysis. We extend the GenNet framework to create interpretable neural networks for multiple omics inputs and apply it to a dataset with transcriptomics and methylomics data. We validate the method using four cohorts in the BIOS consortium for the application of predicting age, low-density lipoproteins (LDL) levels and smoking status. Age prediction from methylation or gene expression data has been an active research area popularized by the work of Hannum et al. and Horvath^23,24^. Additionally, it has been shown that these clocks show an asymptote for older participants and strong biological sex differences, making age prediction particularly interesting to study with neural networks^25^. Smoking status and LDL level predictions are well-suited to evaluate the performance, stability and interpretation of the method.

Methylation and gene expression are highly predictive for smoking status and predictive genes are well-documented^26,27^. On the other hand, low lipid lipoprotein cholesterol levels is a complex outcome with both environmental and genetic factors^28^.

To summarize, we develop visible neural networks for multi-omics data and investigate their generalizability and robustness for three different phenotypes by leveraging the multi-cohort setting of the BIOS consortium in a cohort-wise cross validation analysis. Furthermore, we use the flexibility and interpretability of these models to find sex-specific effects, omic-specific information and genes and pathways important for prediction

## Materials & Methods

### BIOS

In this study, multi-omics data gathered by the Biobank-based Integrative Omics Study (BIOS) consortium was used to predict smoking status, age and low-density lipoprotein levels. Specifically, we used transcriptome and methylome data from BIOS four largest cohorts; LifeLines (LL), Leiden Longevity Study (LLS), Netherlands Twin Register (NTR), and Rotterdam Study (RS). All cohorts within the BIOS consortium followed the same procedure in gathering and processing data. For each participant, transcriptome and the methylome were measured in whole blood samples taken from the same visit. DNA methylation was profiled according to the manufacturer’s protocol using the Infinium Illumina HumanMethylation 450k arrays, while blood was first depleted from globin transcripts for RNA sequencing. A detailed description of all data generation and preprocessing steps for the RNA sequencing and DNA methylation data can be found in Zhernakova et al. (2017)^29^ and Bonder et al. (2017)^30^. Using the BBMRI-NL’s Integrative Omics analysis platform^31^, all individuals that had both RNA-seq and methylation data (β-value) available were selected, resulting in a dataset with 2940 individuals. Y-chromosomal data was excluded, X-chromosomal and autosomal measurements were included. Finally, RNA-seq expression data was filtered using an expression inclusion criterion of one count per million on average across all samples or higher^9^. An overview of the characteristics for each cohort can be found in Table 1.

**Table 1.** Main characteristics for all cohorts used in this study. Note the age differences between the cohorts; participants of the Netherlands Twin Register were on average 29 years younger than the participants of the Rotterdam Study. *Former smokers were excluded in this study. CI; confidence interval.

|  | Rotterdam Study | LifeLines | Leiden Longevity Study | Netherlands Twin Register | Total of all cohorts |
| --- | --- | --- | --- | --- | --- |
| Abbreviation | RS | LL | LLS | NTR |  |
| Individuals | 693 | 727 | 646 | 874 | 2940 |
| Sex, male \| female | 397 \| 296 | 421 \| 306 | 340 \| 306 | 577 \| 297 | 1735 \| 1205 |
| Smokers*, current \| never | 75 \| 231 | 107 \| 337 | 75 \| 184 | 155 \| 500 | 412 \| 1252 |
| Age [years], mean + 95% CI | 67.6<br>(67.1 - 68.0) | 45.4<br>(44.4 - 46.3) | 58.8<br>(58.3 - 59.3) | 38.3<br>(37.3 - 39.3) | 51.4<br>(50.9 - 52.0) |
| LDL [mmol/L], mean + 95% CI | 3.32<br>(3.26 - 3.39) | 3.19<br>(3.12 - 3.25) | 3.36<br>(3.29 - 3.43) | 2.90<br>(2.84 - 2.96) | 3.17<br>(3.14 - 3.2) |

Neural network architectures were created using principles from the GenNet framework^18^. This framework uses prior knowledge (e.g., gene and pathway annotations) to connect input data to the neurons in the next layer of neural network. CpG methylation sites were annotated using GREAT^32^ (Genomic Regions Enrichment of Annotations Tool) and connected to the closest gene based on genomic distance (in base pairs) resulting in 17,283 gene annotations for 481,388 methylation sites. These gene annotations were intersected with the 14,248 remaining gene expression measurements left after preprocessing, resulting in an overlap of 10,404 genes between both omic types. This set of overlapping genes was used in all analyses. The methylation gene layer was built using these genes and their corresponding 324,295 CpGs. For the creation of pathway layers the set of overlapping genes was grouped into KEGG’s functional pathways^33^ from ConsensusPathDB^34^. Out of the 10,404 genes, 4813 genes were annotated for at least one pathway.

**Table 1.** Performance for cohort-wise cross validation, mean with 95% confidince interval over 10 runs. The area under the curve is reported for the classification task (smoking status prediction) and the root mean squared error (RMSE) for the regression tasks; predicting age and LDL levels. ME; Methylation, GE; Gene expression, ME+GE, both methylation and gene expression as an input for the neural network. RS; Rotterdam study, LL; LifeLines, NTR; Netherlands Twin Register, LLS; Leiden Longevity Study. See Supplementary Table 1 for the performance of out-of-the-box scikit-learn implementations for each omic.^*^The name of the test cohort is used to denote the fold. Thus, for the first fold, RS, was used for testing and LL+LLS+NTR were used for training and validation (75% training, 25% validaton).

| Phenotype | Network type | AUC validation cohorts |  |  |  | AUC test cohort |  |  |  | Mean test AUC over all cohorts |
| --- | --- | --- | --- | --- | --- | --- | --- | --- | --- | --- |
|  |  | RS* | LL* | LLS* | NTR* | RS | LL | LLS | NTR |  |
| Smoking | ME+GE | 0.92<br>(0.90 - 0.94) | 0.93<br>(0.92 - 0.93) | 0.91<br>(0.90 - 0.92) | 0.93<br>(0.91 - 0.94) | 0.98<br>(0.98 - 0.98) | 0.92<br>(0.92 - 0.93) | 0.95<br>(0.95 - 0.96) | 0.91<br>(0.89 - 0.92) | **0.95**<br>(0.90 - 1.00) |
|  | ME | 0.93<br>(0.92 - 0.94) | 0.94<br>(0.94 - 0.95) | 0.95<br>(0.94 - 0.95) | 0.93<br>(0.93 - 0.94) | 0.97<br>(0.97 - 0.98) | 0.94<br>(0.93 - 0.94) | 0.96<br>(0.95 - 0.96) | 0.95<br>(0.95 - 0.96) | **0.95**<br>(0.93 - 0.98) |
|  | GE | 0.83<br>(0.83 - 0.84) | 0.82<br>(0.82 - 0.82) | 0.83<br>(0.83 - 0.83) | 0.87<br>(0.86 - 0.87) | 0.87<br>(0.87 - 0.88) | 0.85<br>(0.85 - 0.85) | 0.87<br>(0.87 - 0.88) | 0.80<br>(0.80 - 0.80) | **0.85**<br>(0.80 - 0.90) |
| Phenotype | Network type | RMSE validation cohorts |  |  |  | RMSE test cohorts |  |  |  | Mean test RSME over all cohorts |
|  |  | RS* | LL* | LLS* | NTR* | RS | LL | LLS | NTR |  |
| Age | ME+GE | 3.85<br>(3.72 - 3.98) | 4.13<br>(4.03 - 4.23) | 3.88<br>(3.78 - 3.98) | 4.12<br>(3.99 - 4.25) | 5.44<br>(5.23 - 5.65) | 6.04<br>(5.76 - 6.31) | 4.20<br>(4.11 - 4.29) | 6.33<br>(5.87 - 6.8) | **5.16**<br>(3.97 - 6.35 ) |
|  | ME | 14.48<br>(13.04 - 15.92) | 8.23<br>(7.26 - 9.19) | 18.06<br>(15.01 - 21.11) | 11.28<br>(9.90 - 12.66) | 15.69<br>(12.37 - 19.0) | 8.14<br>(6.50 - 9.79) | 13.17<br>(9.13 - 17.2) | 22.66<br>(19.67 - 25.65) | **15.00**<br>(4.31 - 25.69 ) |
|  | GE | 6.79<br>(6.77 - 6.81) | 6.52<br>(6.51 - 6.53) | 6.40<br>(6.38 - 6.41) | 6.16<br>(6.14 - 6.17) | 9.03<br>(8.80 - 9.26) | 12.23<br>(12.04 - 12.41) | 6.87<br>(6.80 - 6.95) | 17.77<br>(17.51 - 18.03) | **11.45**<br>(4.21 - 18.68 ) |
| LDL | ME+GE | 0.88<br>(0.87 - 0.88) | 0.88<br>(0.88 - 0.88) | 0.85<br>(0.84 - 0.86) | 0.92<br>(0.91 - 0.92) | 0.92<br>(0.92 - 0.93) | 0.89<br>(0.89 - 0.90) | 0.93<br>(0.93 - 0.94) | 0.93<br>(0.91 - 0.95) | **0.93**<br>(0.88 - 0.99) |
|  | ME | 1.18<br>(1.12 - 1.25) | 1.14<br>(1.06 - 1.22) | 1.10<br>(1.02 - 1.18) | 1.12<br>(1.08 - 1.16) | 1.29<br>(1.15 - 1.44) | 1.17<br>(1.07 - 1.26) | 1.10<br>(1.01 - 1.19) | 1.23<br>(1.15 - 1.30) | **1.27**<br>(0.8 - 1.75) |
|  | GE | 0.88<br>(0.88 - 0.88) | 0.88<br>(0.88 - 0.89) | 0.85<br>(0.85 - 0.86) | 0.90<br>(0.90 - 0.91) | 0.94<br>(0.93 - 0.94) | 0.90<br>(0.89 - 0.90) | 0.93<br>(0.93 - 0.93) | 1.02<br>(0.98 - 1.07) | **0.98**<br>(0.79 - 1.17) |

The gene expression network (GE network, Figure 1a) is the simplest network and consists of the gene expression input connected straight to the output node similar as in LASSO regression. The methylation network (ME network, Figure 1b) which consists of the input methylation data, a gene layer with neurons representing gene methylation made and an output node. The methylation and gene expression network (ME + GE network, Figure 1c) combines both networks. In a similar way as in the ME network, CpGs are fed to the first layer of the network and reduced to one node per gene using gene annotations. In contrast to most other methods, gene expression is not concatenated to the input but used as a separate input in the gene level of the network. In this layer, gene expression is combined with the neurons representing genes by methylation. Finally, a single node was used to predict the target phenotype.

**Figure 1.**
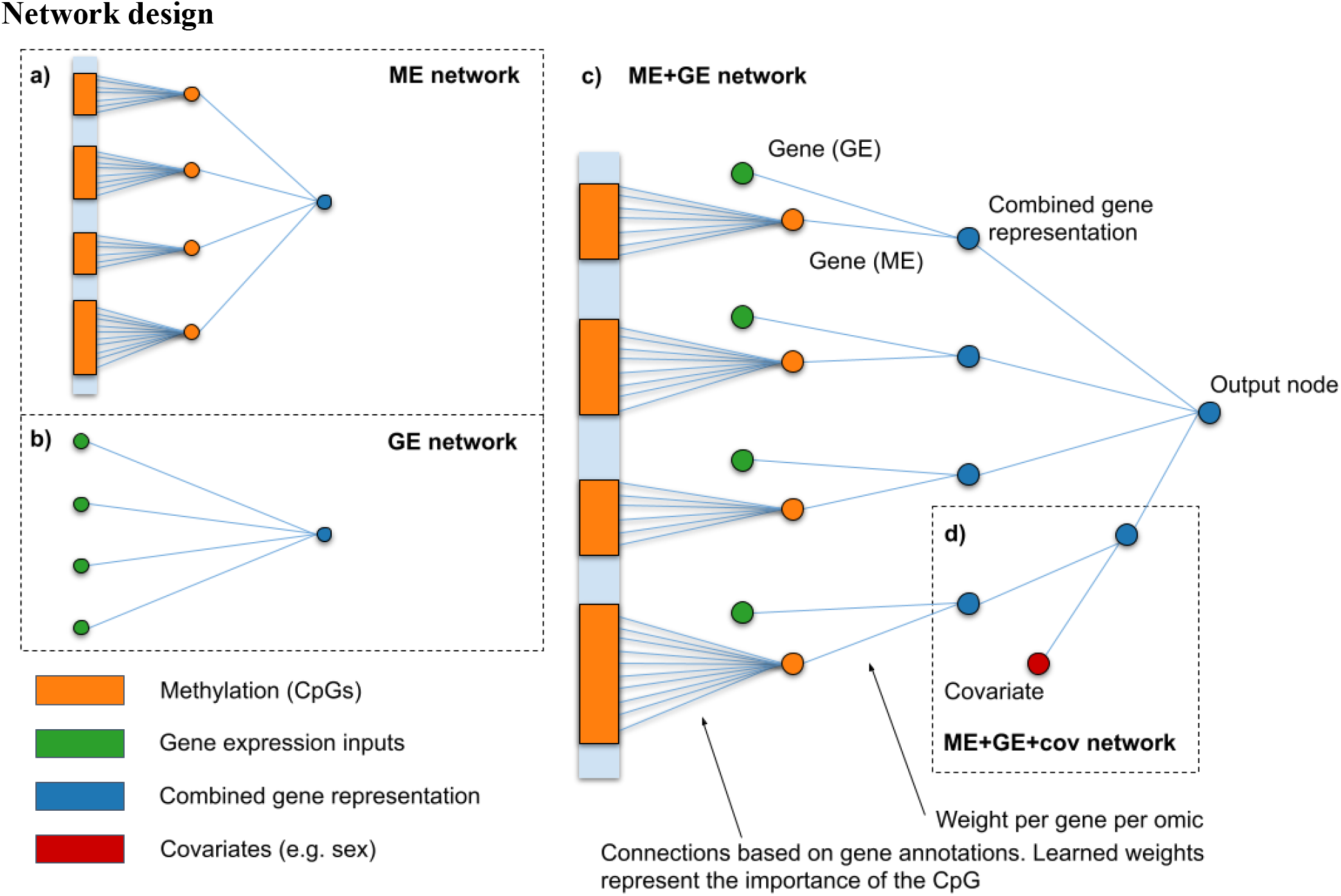
Schematic overview of the neural network architectures used in this study. In the ME network (**a**), DNA methylation data (CpGs) are grouped and connected using gene annotations. The resulting 10,404 gene nodes are directly connected to the output node. Combining the ME network and the the GE network (**b**) for gene expression, results in the ME+GE network (**c**). In the ME+GE network each gene has a node per omic and a combined gene representation. Design (**d**) adds a covariate input to the combined gene representation for each gene. This allows the ME+GE network to model gene-specific effects for the covariate. A schematic overview of the pathway network and a similar fully connected network can be found in Supplementary Figure 1.

The activation function transforms the output signal for each neuron. For classification tasks, such as predicting if an individual smokes or not, a sigmoid activation function was used to scale the output to the range [0, 1] in the last neuron. Arctanh activation functions were used for all other layers to introduce non-linearities, increasing the modelling capabilities of the network. For regression tasks, such as predicting continues traits such as age and LDL levels, ReLu activation functions with output range [0, ∞) were used for all layers. For a better initialization of the network, the bias of the last neuron was set to the mean value of the predicted outcome in the training set.

### Deeper networks

For more complex modelling of the interactions between expression, methylation and phenotypes, we also evaluated deeper networks (Supplementary Figure 1). First, using KEGG’s functional pathways^33,34^ as prior knowledge, three hierarchical pathway layers were created. The first layer groups genes into 321 functional pathways such as: *insulin secretion, thyroid hormone synthesis* and *PPAR signaling pathway*. The forementioned pathways are all part of the endocrine system group which, in turn, is a subgroup of organismal systems. The mid and global-level pathway layers were created adopting this hierarchical structure, consisting of 44 and 6 groups, respectively. Each pathway is represented by its own neuron resulting in three layers with 321, 44 and 6 nodes each. Not all genes were annotated by the KEGG functional pathway annotations, 5591 genes did not receive a functional pathway annotation. To ensure connectivity to the output for all genes, connections that skip the pathway layers (skip connections) were added from each gene to the output node.

Additionally, a deeper network was constructed without any additional prior biological knowledge to compare with the KEGG pathway network. Similarly, the ME+GE network served as a basis for this network and three densely connected layers, 321, 44 and 6 nodes each, were added between the gene layer and the output node. The resulting network has thus the same number of neurons as the KEGG pathway network, but has fully connected layers instead of layers based on KEGG pathway information.

### Training and evaluation

The neural networks were evaluated in a cohort-wise cross validation setup as shown in Figure 2 to assess the generalizability of the models across cohorts. Each fold, one cohort is held out as test set, while the three other cohorts were used for training and validation (leave-one-out method). From these three cohorts 75% of the individuals were randomly selected for the training set while the remaining 25% was used in the validation set to tune the hyperparameters. For all methods the same combinations of hyperparameters were tuned on the validation set. Combinations included learning rates of [0.01, 0.001, 0.005, 0.0001] and L1 penalty on the weights of the combined gene and/or methylation gene layer of [0.01, 0.001, 0.0001]. A higher L1 penalty increases the cost for the network to include more contributors to predict the outcome. The L1 penalty thus enforces sparsity over the weights, so that most inputs get assigned a (near) zero weight while important inputs still get assigned a high weight. This L1 regularization on the weights helps preventing overfitting and increases interpretability.

**Figure 2.**
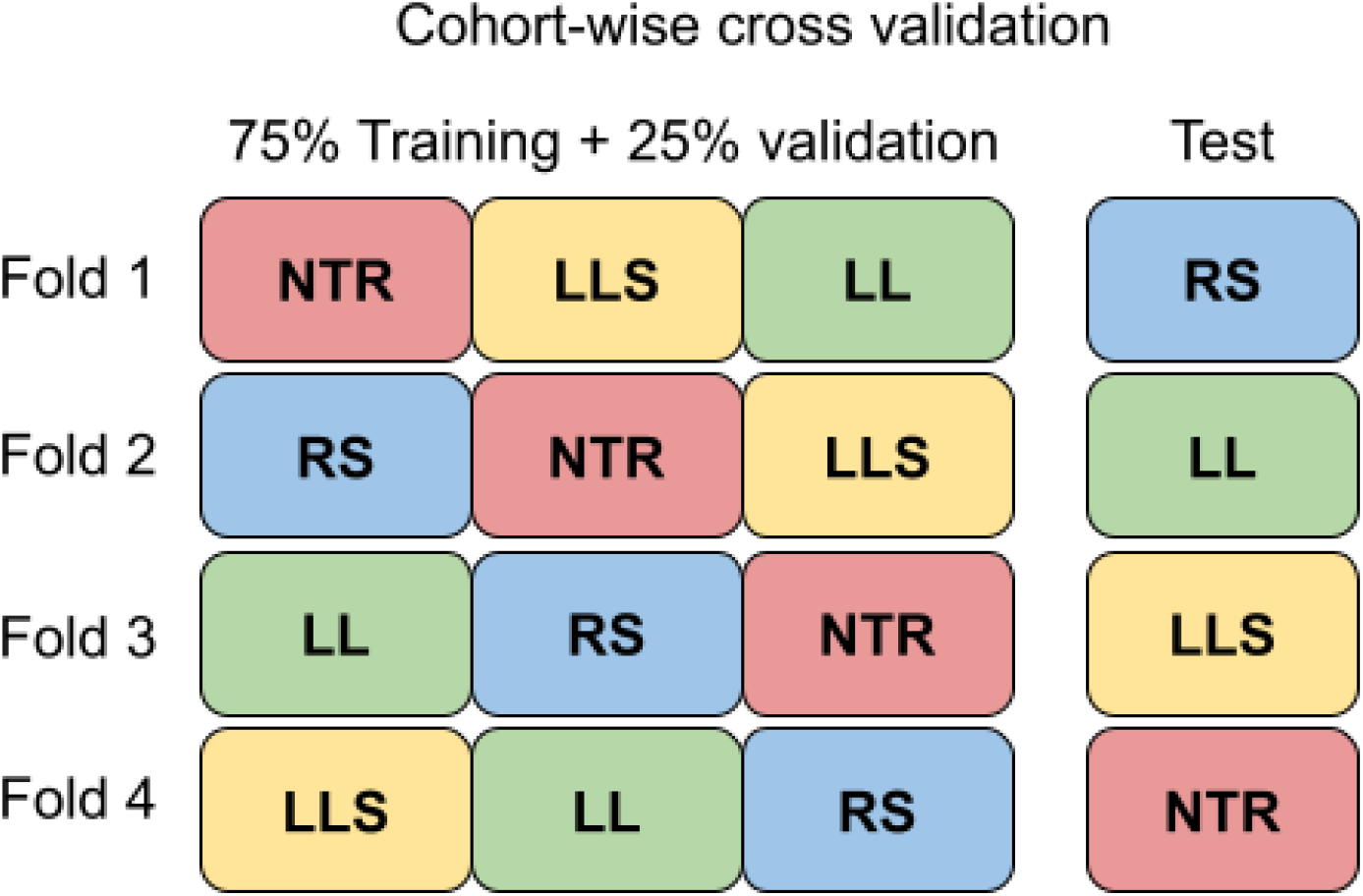
Cohort-wise cross-validation. For each fold three cohorts were used to train and validate the hyperparameters of the model (75% training, 25% validation). The remaining, left-out cohort served as an independent test set and the average performance over the test cohorts was reported. The cohort-wise cross validation was done for each phenotype (smoking, LDL and age prediction). Abbreviations: Netherlands Twin Register (NTR), Leiden Longevity Study (LLS), LifeLines (LL), Rotterdam Study (RS).

The mean squared error (MSE) was used as a loss function to optimize for regression tasks. For classification tasks, weighted binary cross entropy, with a weight inverse to the ratio of the class imbalance, was used as a loss function. The loss function quantifies the difference between current outcome and the true label and is optimized during training. The performance of the resulting network is evaluated using the area under the receiver operating curve (AUC) for classification tasks, and the root mean squared error (RMSE) and explained variance for regression tasks (explained variance can be found in the Supplementary Materials). For each fold the hyperparameters of the best performing model in the validation set were selected to evaluate on the test cohort. Since neural networks use stochastic processes that can influence the outcome, we trained the network with the best hyperparameters ten times with a different random seed to investigate its stability.

### Additional analyses

Neural networks are flexible methods and with the inclusion of prior biological knowledge different architectures can be explored to provide more insight into the interaction between omics types, contribution of covariates and gene-specific contribution of covariates. For each of these analyses we made small changes to the ME+GE networks.

#### Omic-specific information

Gene expression and methylation data contain redundant information with respect to each other. However, not all information that is present in the one may be present in the other data type. To evaluate the independent contribution of each omics to the prediction we add a L1 penalty for one omics type in the model. This introduces a trade-off for the neural network: the gain in performance for including information of the penalized omic (i.e., RNA expression of a single gene or the methylation representation of a single gene) must outweigh its penalty. If the model uses only the non-penalized omic type without loss of prediction performance, it is likely that there was no omic-specific information. However, if the model still decides to use parts of the penalized omics data, this information is most likely unique to the penalized omic type and was therefore required for prediction.

#### Covariate-gene interaction

Including covariates in the model, for example sex and age for smoking, can improve performance and interpretation. Commonly, the covariates are included as an extra layer at the end. However, by adding a covariate for each gene, more specific information in how a covariate affects a single gene can be obtained (see Figure 1d). For each phenotype we tested both, a model with covariates in the last layer and a model with covariates for each gene.

#### Subtyping with activation patterns

In contrast to fully-connected neural networks, the visible neural network architectures used in this study are constructed based on prior biological knowledge can be interpretated by inspecting the weights of the incoming and outgoing connections. The strength of the weights (e.g., between CpGs and genes, expression and genes, genes and pathways), all express the importance of these biological elements for the predicted outcome. The weights of a neural network are a result of an optimization over the population it was trained on, and are thus a result of the population characteristics of the training set. However, neural networks may learn different patterns for the same outcome. By inspecting the weights general information is learned about the importance of each element but this does not show differences between groups or individuals. Based on differences between individuals, some neurons can activate for a certain group of individuals, while it does not for others. To gain an overview of the different patterns that are learned by the network we applied principal component analysis^35^ (PCA) over all the activations for all (gene-level) nodes for each individual. In this PCA, individual-level differences may cluster and provide groups of individuals for which the neural network used a similar activation pattern.

## Results

### Cohort-wise cross validation

An overview of the performance for each cohort and for the three different architectures can be found in Table 2. It shows the mean predictive performance and standard deviation for each fold for ten networks trained with the same hyperparameters but with different random seeds. The corresponding hyperparameters, chosen on the best performance in the validation set, can be found in Supplementary Table 2.

#### Predicting smoking status

Both gene expression and methylation were highly predictive for smoking status in all folds. The best performance was achieved by the ME+GE network, thus with both methylation and gene expression input, in the fold with the Rotterdam Study as test cohort (all other cohorts were used for training and validation). In this fold, the network achieved a near perfect classification with area under the receiver operating curve (AUC) of 0.98 (95% confidence interval, 0.98 - 0.98). Over all folds, the ME networks and ME+GE networks performed best with a mean AUC of 0.95 (95% CI, 0.93 - 0.98) and 0.95 (95% CI, 0.90 - 1.00) respectively. The GE network, based solely on gene expression input, performed substantially worse with a mean AUC of 0. 0.85 (95% CI, 0.80 - 0.90). Surprisingly, the mean test performance over all folds for the ME+GE network was lower for deeper networks with three fully connected layers, achieving a mean AUC of 0.91 (95% CI, 0.85 - 0.96) (see Supplementary Table 3). In general, each fold obtained good predictive performance for predicting smoking status, the GE network in the fold with NTR as test cohort achieved the worst overall predictive performance with a mean AUC of 0.80 (95% CI, 0.80 - 0.80).

The ME+GE networks exhibit a stable performance, with small confidence intervals for the area under the curve and standard deviations not exceeding 0.03. However, there may be significant variations in the underlying weights due to stochastic processes used for network initialization and training, resulting in different starting points and optimization paths for all weights across runs. As the weights within a neural network operate relative to each other and cannot be directly compared between networks, we compared the relative contribution of each gene instead. Figure 3 demonstrates that certain genes are consistently utilized by the network to differentiate between current smokers and non-smokers across all folds, although there can be notable differences in the percentage of total weight each gene holds. In each fold, *GPR15* is the most or second most predictive gene for smoking status, its signal is mainly driven by gene expression as visualized in Figure 3. Specifically, 79.8 ± 33.3% (mean and standard deviation over all folds) of the weights that drive the signal for this gene are from the gene expression input. The next gene, *AHRR*, is important for prediction in three out of four cohorts. This signal is driven by both gene expression (44.3%) as well as methylation (55.6%). Other consistently highly predictive genes (i.e., genes with a weight contribution higher than 1% in three out of four cohorts) are *SEMA6B, PID1, LRRN3, P2RY6, CDKN1C, CLEC10A* and *KCNQ1*. (See Supplementary Table 4 for more details). All these consistently highly predictive genes were found before in association studies for smoking in gene expression and methylation^36–38^. A graphical overview of important pathways for smoking prediction can be found in Supplementary Figure 2.

**Figure 3.**
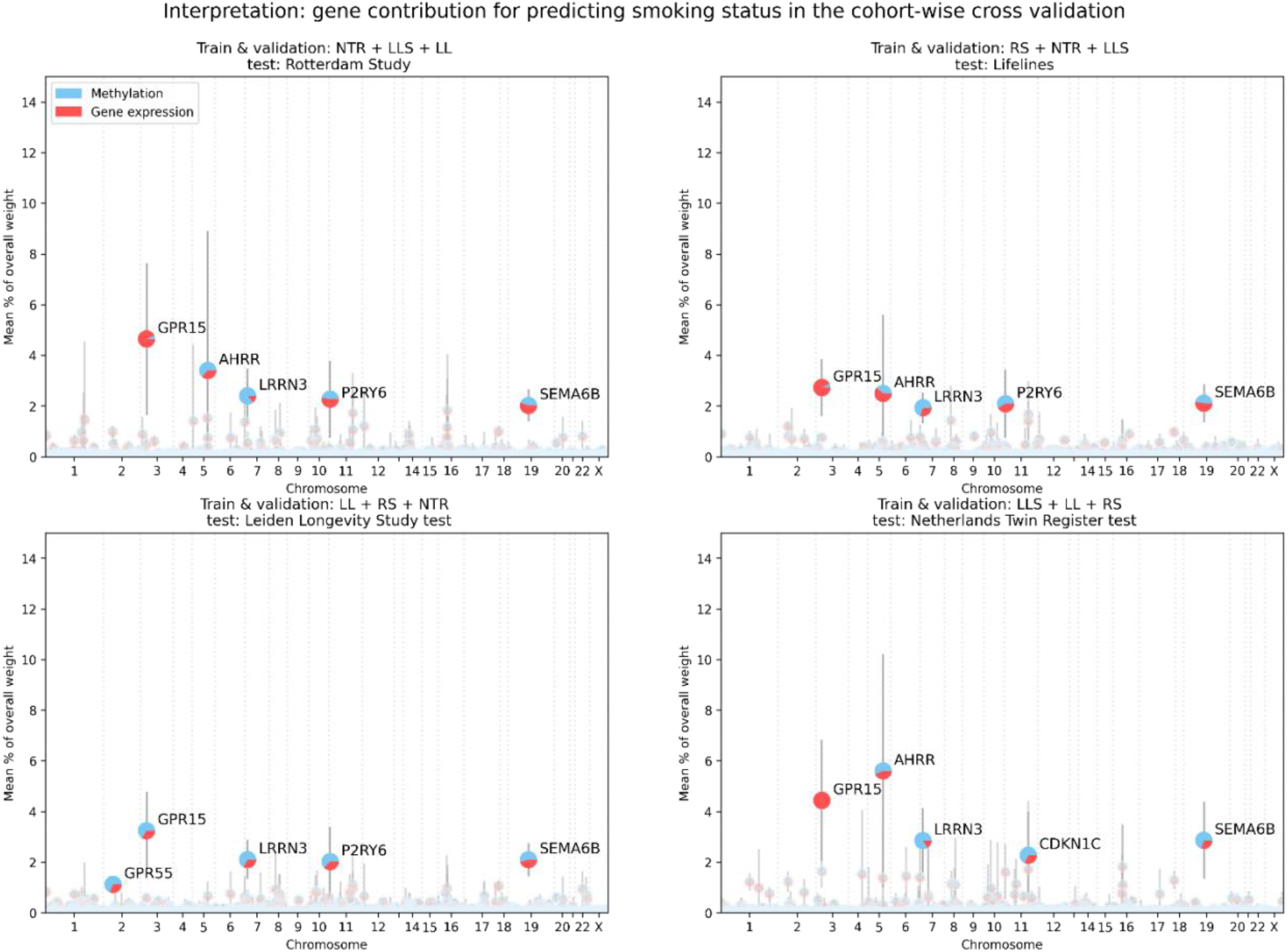
Overview of the important genes for predicting smoking status for each fold. Contribution was measured in percentage of the total weight assigned to each gene. For each gene, the pie chart shows the contribution of methylation and gene expression. Error bars indicate the standard deviation over ten runs for the exact same network trained with the same hyperparameters.

To investigate the interplay between the omic types, two additional analyses were conducted where either gene expression or methylation gene representations were penalized (see Supplementary Figures 3,4 and 5). Without penalization the weights for gene expression and methylation were nearly equally divided after training. Weights connected to gene expression input occupied 51.6 ± 1.3% of the weights over all the ME+GE networks, the remainder used for methylation. In these experiments, we found that an omic specific L1 penalty of 0.01 for gene expression reduced the contribution of the weights associated with gene expression to 0.69 ± 1.16% while a similar threshold reduced the weights associated with methylation to 2.56 ± 1.73%. A more severe omic specific L1 threshold of 0.001 for methylation reduced the use of methylation in the top genes nearly completely, only for *LRNN3* methylation input is still used in the second and third fold with (respectively ∼41% and 16% of the weights for this gene). However, with the same threshold gene expression inputs are responsible for 15% of the weights for AHRR in the first fold, nearly 29% of the *GPR15* weights in the second fold and 39% of *RER1* in the fourth fold (see **Error! Reference source not found**. and 4). Interestingly, the importance of AHRR was severely impacted by the methylation penalty, its gene expression was barely used to predict smoking status when methylation was penalized.

#### Predicting age

Networks trained with both methylation and gene expression data (ME+GE) achieved a mean error of 5.16 (95% CI, 3.97 - 6.35) years over all folds for age prediction (see Table 1). Between folds, there were large differences in performance for predicting age. Most notably, networks did not generalize well in folds that have either the Rotterdam Study (ranging between 52 to 80 years) or the Leiden Longevity Study (ranging between 30 to 79 years) as test cohort, the two cohorts with the oldest population. For these cohorts, the explained variance in the test set was substantially lower than in the validation set: Rotterdam study test 0.40 (95% CI, 0.37 - 0.43), 0.94 (95% CI 0.93 - 0.94) validation, Leiden Longevity Study test 0.95 (95% CI, 0.95 - 0.95), 0.61 (95% CI, 0.60 - 0.63) validation. Aside from being older, these cohorts also have a smaller spread in age distribution compared to the two other cohorts (See Figure 4a and Supplementary Figure 6). The Netherlands Twin Register cohort ranges between roughly 18 and 80 years old while individuals from the Lifelines cohort were between 18 and 81 years old.

**Figure 4.**
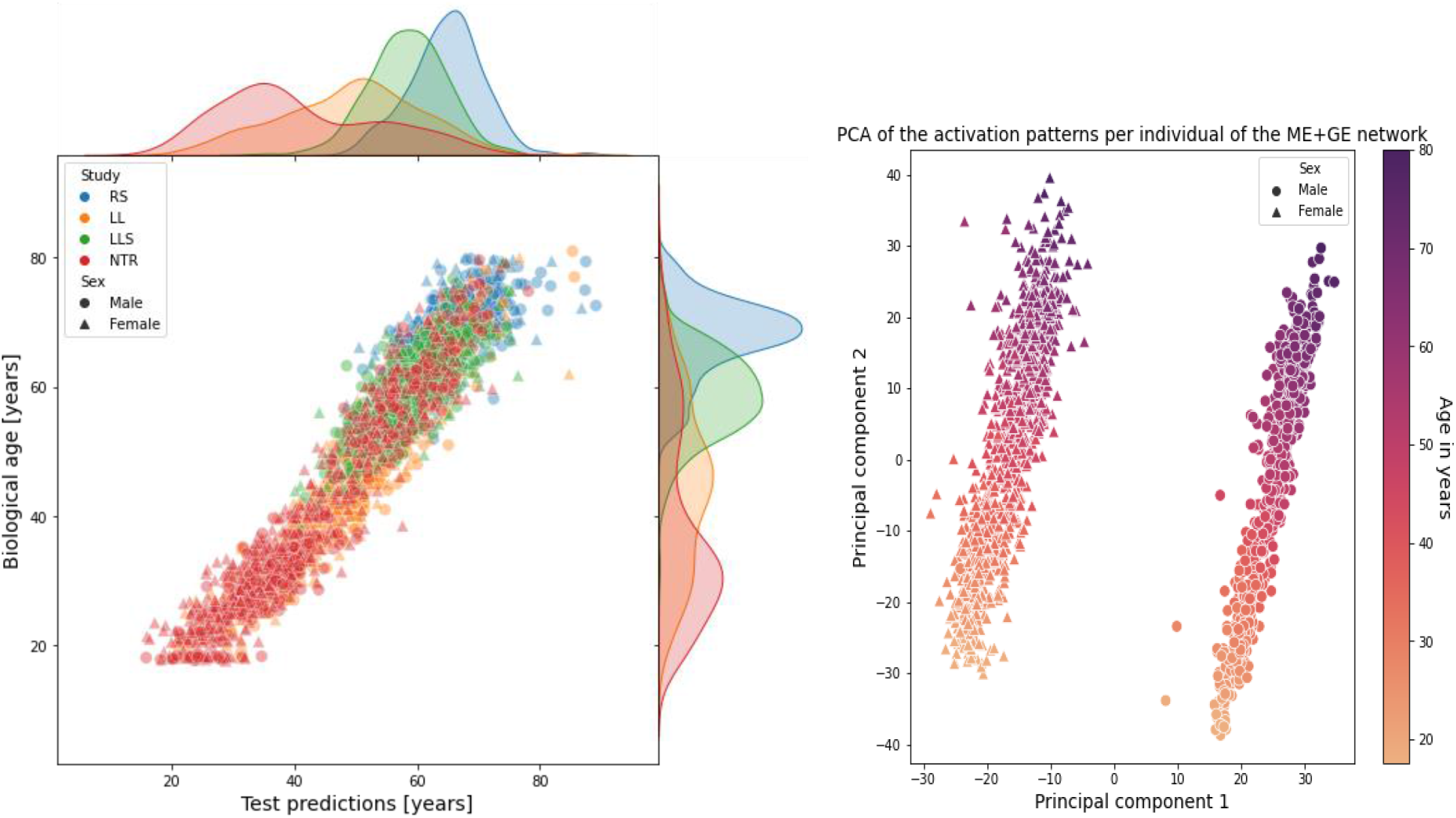
a) Test predictions for the ME+GE network for all folds (each cohort) with corresponding distributions (See Supplementary 11 and 12 for the GE and ME networks). b) Activation of the ME+GE trained for age prediction. A principal component analysis clearly shows two distinct activation patterns corresponding to the different sexes. Principal component 1 is related to the sex differences, principal component 2 to the age of the participants.

Differences between omics and network types were also larger for age prediction than for smoking status prediction. The ME+GE network consistently outperformed the single-omic networks with substantial margins: the mean explained variance over all folds was 0.72 (95% CI, 0.36 - 1.07) for the ME+GE network, 0.30 (95% CI, -0.26 - 0.86) for gene expression, while the ME networks did not find any predictive pattern that translated to the test cohort. Training and validation performance was generally poor for the ME network, and although the GE network obtained good validation performance in terms of explained variance for each fold, this did not translate in folds with the Rotterdam study and Leiden Longevity Study as test cohorts.

Interpretation of the ME+GE network revealed that many genes had a small contribution for age prediction (see Supplementary Figure 6). The neural network found a more multifactorial solution for age prediction than for smoking, the most important gene over all folds only occupied 0.68% of all weights for predicting age compared to 3.76% for smoking. The most predictive genes with a weight contribution higher than 0.30% of the total weight in three out of the four folds were *COL11A2, AFAP1, OTUD7A, PTPRN2, ADARB2* and *CD34* (Supplementary Table 5). These most predictive genes were not part of Hannum et al. and Horvath’s epigenetic clocks^23,24^.

The first principal components of the activation patterns of the ME+GE network revealed distinct activation patterns for the different sexes with a gradient in each cluster (see 4b). Although, there is no significant difference in the absolute error between the sexes (Wilcoxon rank-sum, p-value of 0.98, Supplementary Figure 7,8), the first principal component clusters perfectly for males and females while the second principal component is strongly related with age. Additional experiments showed that the clustering of the sexes is mainly driven by genes on the X chromosome (see Supplementary Figure 9). Including sex as a covariate in the last layer of the model did not improve the performance of the model (mean RMSE over all folds of 7.31 [95% CI, 2.89 - 11.73]). Including sex information to each gene also did not lead to a better performance (mean RMSE over all folds of 10.64 [95% CI, 4.12 - 17.15]). However, inspecting the weights between the covariate and the genes for the best performing network revealed strong sex-specific weights for, among others: *KLF13, ANO9* and *HECA* (for more details see Supplementary Figure 10). For these genes the network needed strong weights to model sex-specific effects for age prediction. After applying an omic-specific L1 penalty for methylation of 0.01, the network only used the methylation input for gene *NEDD1* in the second fold with nearly 33% of the weight contribution for this gene from methylation, while in the third fold *MAD1L1* had a methylation contribution of 23% (see Supplementary Figure 14). With the same threshold for penalizing gene expression inputs, *DNAJB6* had the largest gene expression use with 31% of the weight for this gene assigned to gene expression input (Supplementary Figure 15). The deeper neural network architectures quickly overfitted, reaching high performance on the training data which did not generalize to the validation and test set. These networks were consistently outperformed by the ME+GE network (Supplementary Table 5). The best performing network build with KEGG pathway information had the pathway: “environmental information processing” as the most predictive global pathway because of high contributions of membrane transport (ABC transporters), signal transduction, and signaling molecules and interaction (see Supplementary Figure 16).

#### Predicting low-density lipoproteins levels

ME+GE and GE networks explained up to 17% of the phenotypic variance in the validation set but these networks only generalized in the second fold to an explained variance of 0.07 (95% CI, 0.05 - 0.08) for the ME+GE network and 0.04 (95% CI, 0.04 - 0.05) for the GE network in the Lifelines test cohort (see **Error! Reference source not found**.). In this fold, the largest gene, *FAM53A* only occupied 0.052% of the total weight (Supplementary Figure 17). The weights for all genes in the ME+GE network were small and evenly spread, indicating that the network did not find individual genes with a strong effect for predicting low-density lipoproteins levels. Additional layers, be it pathways or densely connected layers, did not improve predictive performance.

## Discussion

In this paper we evaluated the performance, interpretability and stability of visible neural networks for single and multi-omics data. Interpretability was achieved by embedding prior biological knowledge such as gene and pathway annotations in the neural network architecture. We applied these models to predict smoking status, age and low-density lipoprotein levels in a cohort-wise cross validation using methylation and gene expression data.

For smoking, single omic networks and multi-omic networks performed consistently high across all cohorts for predicting smoking status. Predicting smoking status is a relatively simple task, since smoking is a powerful inducer of DNA methylation and gene expression alterations^39^. This is also reflected by the mean AUC of 0.95 over all folds that the ME+GE and ME networks achieved. It is slightly better than the performance of Maas et al. who reported an AUC of 0.90 in an external dataset with a weighted combination of just thirteen CpGs. Inspection of the weights of the ME+GE network revealed *GPR15, AHRR* and *LRRN3* as most important genes for prediction, which is consistent with existing literature^26,27,39,40^. In the ME+GE network the contribution of both omics types was nearly equal (in terms of weights), while the gene expression-based network by itself was less predictive than the methylation-based networks. Applying an omic-specific penalty for methylation input showed that the ME+GE network needed some methylation input to achieve similar performance with expression information.

For predicting age, the ME+GE network outperformed the single ME or GE networks. The performance of this network in the test cohorts varied between a R^2^ of 0.40 (95% CI, 0.37 - 0.43) and 0.91 (95% CI, 0.90 - 0.92). This difference in performance is probably caused by the different distributions in age in the cohorts, depending on the cohorts in the training set the networks are shown less examples of older or younger individuals. Similar effect were also seen in traditional methods^9^. Based on the predictive performance shown in Table 2 one could conclude that for age prediction, usage of the two omics types increased stability and performance for these type of neural networks compared to the single omic networks. Additionally, we have evaluated whether the network used sex-information in the decision process for age prediction. The first principal component of the activations of the neural network showed a perfect separation between the sexes, mostly caused by genes on the X-chromosome, while the second principal component had a clear correlation with age. Owing to the shallowness of the networks, the activation pattern will therefore closely resemble the underlying data, especially if it has some relation with the outcome. For deeper networks a PCA on the activation may reveal more detailed information (such as different patients subtypes or mediating factors) since the network applies more complex transformations to the data. The inclusion of genes on the X-chromosome allowed the network thus to separate between the sexes but it did not have the capacity to model different effects independently from the input for each sex. To help the model to find a sex-specific effect we modified the network with sex information as an extra input to each gene node. After, training the network found the strongest sex-specific gene effects for *KL13, ANO9* and *HECA*. However, this addition to the network architecture did not improve performance.

An earlier EWAS in only the Rotterdam study did not find significant associations between DNA methylation in blood and low-density lipoproteins cholesterol^41^. Another EWAS using BIOS data found only three significant associations, demonstrating that there is a very weak relation between methylation and LDL measurements from blood which makes the prediction task more complex^42^. The neural networks did find patterns in the training set that were also found in the validation set (up to an R^2^ of 0.17 [95% CI, 0.16 - 0.18]) but this pattern did not generalize to the test cohorts with the exception of the Lifeline cohort. In this cohort the method achieved an R^2^ of 0.07 (95% CI, 0.05 - 0.08) in the test set, substantially lower than the performance of the validation set 0.13 (95% CI, 0.12 - 0.14). suggesting that the model had trouble generalizing to data from an unseen cohort. Overall, the low prediction performance might also indicates that the studied omic-data (gene expression and methylation from blood) might not contain enough information to accurately predict LDL-levels.

In general, we found that including multiple omics inputs in the network improved performance. These multi-omic networks had a more stable performance and generalized better to the test cohorts. Surprisingly, deeper networks did not lead to better performance. Generally, one would expect deeper networks to perform better since they can model more complex interactions. Thus, it is possible that the optimal hyperparameter values for deeper networks lie outside the considered hyperparameter range or that more training examples are required to train these deeper networks. Interpreting the ME+GE networks revealed well-known genes such as *GPR15* and *AHRR* for smoking that validate the results. However we also saw that the interpretation can vary between different random initializations and it is therefore recommended to train networks with different random seeds for a more complete overview of important predictors. As for all prediction models, it is important to consider that predictive genes and pathways found are not necessarily causal genes and pathways as effects can be mediated. However, these genes and pathways do provide insight in the decision process of the neural network and may be used in follow-up.

For good interpretation, proper regularization is important as it forces the network to use the most predictive input features. For example, an L1 penalty on the weights will force the network to learn sparse weights, resulting in a less complex model. In the absence of an L1 penalty on the weights, the network has more freedom to choose its weights. This does not necessarily harm performance, but may harm interpretability. In this work we use the L1 penalty to regularize the network, but other regularization methods could have been chosen. For example dropout^43^, this method drives the network to find a more stable solution by deactivating random sets of neurons during training. Another important factor for interpretation in visible neural networks is the quality of the prior knowledge used in creation. In this study, the annotations for the CpG sites were based on genomic distance. Potential improvements could come from using tissue specific and functional annotation databases such as ENCODE^44^

## Conclusion

We believe that visible neural networks have great potential for genomic applications, especially for multi-omics integration. These interpretable neural networks can combine multi-omics data elegantly in a single prediction model and provide the importance of each gene, pathway and omic input for prediction. Additionally, we found that using multi-omic networks generally improved performance, stability and generalizability compared to interpretable single omic networks.

## Data availability

BIOS datasets are available from the European Genome-Phenome Archive by accession number EGAS00001001077 (https://ega-archive.org/studies/EGAS00001001077). Alternative options to access the data are available through the BIOS website; https://www.bbmri.nl/acquisition-use-analyze/bios/. All trained networks are available on request.

## Code availability

Code is available on GitHub: https://github.com/ArnovanHilten/GenNet-multi-omic

## Acknowledgements

This work was funded by the Dutch Technology Foundation (STW) through the 2005 Simon Steven Meester grant 2015 to W.J. Niessen. G.V. Roshchupkin supported by the ZonMw Veni grant (Veni, 1936320). The Rotterdam study is supported by the Netherlands Organization for Scientific Research (NWO, 91203014, 175.010.2005.011, 91103012). This research was financially supported by BBMRI-NL, a Research Infrastructure financed by the Dutch government (NWO, numbers 184.021.007 and 184.033.111). Samples used in this study were contributed by LifeLines, the Leiden Longevity Study, the Netherlands Twin Registry (NTR) and the Rotterdam Study. This work was carried out on the Dutch national e-infrastructure with the support of SURF Cooperative. We thank all participants and investigators for their contributions to this study.

## Contributions

A.H., J.v.R. and G.R. conceived and designed the method. A.H. performed experiments and implemented the method. G.R., W.N. and J.v.M supervised the work. Data set generation and quality control of the BIOS datasets was done by the BIOS Consortium, more details can be found at: http://www.bbmri.nl/acquisition-use-analyze/bios/, including details on contributions of all consortium members. A.H., J.v.R, G.R., A.I., W.N., and J.v.M wrote, revised, and approved the paper.

