## Supplementary Materials for "Phenotype prediction using biologically interpretable neural networks on multi-cohort multi-omics data"

#### Supplementary Tables

I. Performance out-of-the-box methods

| Validation set | | | | | |
|---|---|---|---|---|---|
| | | **Rotterdam Study** | **LifeLines** | **Leiden Longevity Study** | **Netherlands Twin Register** |
| **Smoking [AUC]** | ME | 0.71 | 0.71 | 0.69 | 0.75 |
| | GE | 0.80 | 0.83 | 0.80 | 0.82 |
| **Age [expl. var.]** | ME | 0.97 | 0.96 | 0.96 | 0.95 |
| | GE | 0.83 | 0.83 | 0.82 | 0.81 |
| **LDL [expl. var.]** | ME | 0.08 | 0.07 | 0.06 | 0.00 |
| | GE | 0.13 | 0.13 | 0.11 | 0.04 |
| **Test set** | | | | | |
| | | **Rotterdam Study** | **LifeLines** | **Leiden Longevity Study** | **Netherlands Twin Register** |
| **Smoking [AUC]** | ME | 0.77 | 0.77 | 0.70 | 0.73 |
| | GE | 0.93 | 0.80 | 0.88 | 0.77 |
| **Age [expl. var.]** | ME | 0.76 | 0.93 | 0.67 | 0.60 |
| | GE | 0.63 | -0.20 | -0.25 | -0.31 |
| **LDL [expl. var.]** | ME | 0.08 | 0.07 | 0.00 | 0.00 |
| | GE | -0.01 | 0.03 | -0.00 | -0.12 |

*Supplementary Table 1. Performance of out-of-the-box scikit-learn implementations. LogisticRegression with an L1 penalty for predicting smoking status and LASSO for LDL and Age prediction.*

### II. Hyperparameters

| | | | ME+GE | | ME | | GE | | Deep dense | |
|---|---|---|---|---|---|---|---|---|---|---|
| | Fold (by test set) | Abbreviation | L1 | LR | L1 | LR | L1 | LR | L1 | LR |
| Smoking | Rotterdam Study | RS | 0.001 | 0.01 | 0.001 | 0.01 | 0.0001 | 0.001 | 0.001 | 0.001 |
| | LifeLines | LL | 0.001 | 0.01 | 0.001 | 0.01 | 0.0001 | 0.001 | 0.001 | 0.001 |
| | Longevity Study | LLS | 0.001 | 0.01 | 0.001 | 0.01 | 0.0001 | 0.001 | 0.0001 | 0.005 |
| | Netherlands Twin Register | NTR | 0.001 | 0.01 | 0.001 | 0.01 | 0.0001 | 0.001 | 0.001 | 0.001 |
| Age | Rotterdam Study | RS | 0.01 | 0.005 | 0.01 | 0.01 | 0.001 | 0.01 | 0.01 | 0.0001 |
| | LifeLines | LL | 0.01 | 0.005 | 0.01 | 0.01 | 0.001 | 0.01 | 0.001 | 0.001 |
| | Longevity Study | LLS | 0.01 | 0.005 | 0.01 | 0.01 | 0.001 | 0.01 | 0.001 | 0.0001 |
| | Netherlands Twin Register | NTR | 0.01 | 0.005 | 0.01 | 0.01 | 0.001 | 0.01 | 0.01 | 0.1 |
| LDL | Rotterdam Study | RS | 0 | 0.01 | 0.001 | 0.01 | 0.0001 | 0.001 | 0.001 | 0.0001 |
| | LifeLines | LL | 0 | 0.01 | 0.001 | 0.01 | 0.0001 | 0.001 | 0.01 | 0.01 |
| | Longevity Study | LLS | 0 | 0.01 | 0.001 | 0.01 | 0.0001 | 0.001 | 0.001 | 0.000001 |
| | Netherlands Twin Register | NTR | 0 | 0.01 | 0.001 | 0.01 | 0.0001 | 0.001 | 0.001 | 0.000001 |

*Supplementary Table 2. Hyperparameters for the best performance in the validation set. Batch size was in all experiments 64, L1 penalty value (L1) was on the kernel regularization penalty, learning rate (LR) was for the ADAM optimizer.*

### III. Smoking prediction

| Smoking status prediction | | | | | | | | | |
|---|---|---|---|---|---|---|---|---|---|
| **Network type** | **AUC validation cohorts (excluded test cohort)** | | | | **AUC test cohort (test cohort)** | | | | **Mean over all cohorts** |
| **-** | **RS** | **LL** | **LLS** | **NTR** | **RS** | **LL** | **LLS** | **NTR** | |
| **ME+GE** | 0.92 (0.90 - 0.94) | 0.93 (0.92 - 0.93) | 0.91 (0.90 - 0.92) | 0.93 (0.91 - 0.94) | 0.98 (0.98 - 0.98) | 0.92 (0.92 - 0.93) | 0.95 (0.95 - 0.96) | 0.91 (0.89 - 0.92) | **0.95 (0.90 – 1.00)** |
| **ME** | 0.93 (0.92 - 0.94) | 0.94 (0.94 - 0.95) | 0.95 (0.94 - 0.95) | 0.93 (0.93 - 0.94) | 0.97 (0.97 - 0.98) | 0.94 (0.93 - 0.94) | 0.96 (0.95 - 0.96) | 0.95 (0.95 - 0.96) | **0.95 (0.93 - 0.98)** |
| **GE** | 0.83 (0.83 - 0.84) | 0.82 (0.82 - 0.82) | 0.83 (0.83 - 0.83) | 0.87 (0.86 - 0.87) | 0.87 (0.87 - 0.88) | 0.85 (0.85 - 0.85) | 0.87 (0.87 - 0.88) | 0.80 (0.80 - 0.80) | **0.85 (0.80 - 0.90)** |
| **ME+GE (penalized methylation L1 of 0.001)** | 0.88 (0.87 - 0.89) | 0.88 (0.87 - 0.89) | 0.9 (0.9 - 0.91) | 0.91 (0.91 - 0.92) | 0.95 (0.94 - 0.96) | 0.88 (0.88 - 0.89) | 0.94 (0.94 - 0.94) | 0.86 (0.86 - 0.87) | **0.91 (0.83 - 0.99)** |
| **ME+GE (penalized gene expression L1 of 0.001)** | 0.92 (0.90 - 0.94) | 0.92 (0.90 - 0.93) | 0.91 (0.89 - 0.93) | 0.92 (0.91 - 0.93) | 0.98 (0.97 - 0.98) | 0.92 (0.91 - 0.93) | 0.95 (0.95 - 0.96) | 0.89 (0.87 - 0.91) | **0.95 (0.89 - 1.0)** |
| **Deep (5 layers dense)** | 0.83 (0.81 - 0.85) | 0.86 (0.85 - 0.87) | 0.84 (0.83 - 0.85) | 0.89 (0.88 - 0.90) | 0.95 (0.94 - 0.95) | 0.90 (0.89 - 0.90) | 0.89 (0.87 - 0.90) | 0.84 (0.82 - 0.86) | **0.91 (0.85 - 0.96)** |
| **Pathway** | 0.83 (0.82 - 0.84) | 0.83 (0.82 - 0.84) | 0.76 (0.75 - 0.78) | 0.85 (0.84 - 0.85) | 0.89 (0.89 - 0.9) | 0.84 (0.84 - 0.85) | 0.81 (0.78 - 0.84) | 0.8 (0.78 - 0.81) | **0.84 (0.77 - 0.91)** |

*Supplementary Table 3. Results for the base network and variations to the base network. Area under the curve for smoking status prediction for each cohort in a cohort-wise cross validation, mean with 95% confidence interval over 10 runs.*

| Gene | Mean percentage fold 1 | Mean percentage fold 2 | Mean percentage fold 3 | Mean percentage fold 4 | Mean all folds |
|---|---|---|---|---|---|
| GPR15 | 4.64 | 2.73 | 3.24 | 4.44 | 3.76 |
| AHRR | 3.41 | 2.50 | 0.00 | 5.60 | 2.88 |
| LRRN3 | 2.40 | 1.93 | 2.10 | 2.85 | 2.32 |
| SEMA6B | 2.02 | 2.11 | 2.10 | 2.86 | 2.27 |
| P2RY6 | 2.27 | 2.09 | 2.02 | 1.61 | 2.00 |
| CDKN1C | 1.74 | 1.42 | 0.98 | 2.27 | 1.60 |
| KCNQ1 | 1.07 | 1.69 | 1.12 | 1.73 | 1.40 |
| PID1 | 1.00 | 1.20 | 1.08 | 1.24 | 1.13 |
| CLEC10A | 1.00 | 0.98 | 1.07 | 1.30 | 1.09 |

*Supplementary Table 4: Genes with contributions higher than 1% of the total weight for smoking prediction for three out of the four folds.*

#### IV. Biological age prediction

| Age Prediction | | | | | | | | | |
|---|---|---|---|---|---|---|---|---|---|
| | $R^2$ validation cohorts (excluding named cohort) | | | | $R^2$ (test cohort) | | | | Mean over all test cohorts |
| | RS | LL | LLS | NTR | RS | LL | LLS | NTR | |
| **ME+GE** | 0.94 (0.93 - 0.94) | 0.94 (0.94 - 0.94) | 0.95 (0.95 - 0.95) | 0.90 (0.89 - 0.91) | 0.40 (0.37 - 0.43) | 0.88 (0.87 - 0.88) | 0.61 (0.60 - 0.63) | 0.91 (0.90 - 0.92) | **0.72 (0.36 - 1.07)** |
| **ME** | 0.23 (0.10 - 0.36) | 0.81 (0.76 - 0.86) | 0.32 (0.16 - 0.48) | 0.35 (0.23 - 0.48) | -0.30 (-0.87 - 0.27) | 0.75 (0.69 - 0.81) | -0.15 (-0.52 - 0.23) | 0.31 (0.24 - 0.38) | **-0.31 (-2.5 - 1.88)** |
| **GE** | 0.81 (0.81 - 0.81) | 0.85 (0.85 - 0.85) | 0.87 (0.87 - 0.87) | 0.77 (0.77 - 0.78) | -0.08 (-0.09 - -.06) | 0.64 (0.64 - 0.64) | 0.09 (0.09 - 0.09) | 0.55 (0.54 - 0.56) | **0.30 (-0.26 - 0.86)** |
| **ME+GE (L1 ME)** | 0.93 (0.93 - 0.94) | 0.93 (0.93 - 0.94) | 0.95 (0.94 - 0.95) | 0.89 (0.88 - 0.89) | 0.39 (0.34 - 0.44) | 0.87 (0.87 - 0.88) | 0.59 (0.57 - 0.61) | 0.90 (0.89 - 0.9) | **0.69 (0.29 - 1.08)** |
| **ME+GE (L1 GE)** | 0.94 (0.93 - 0.94) | 0.93 (0.93 - 0.94) | 0.95 (0.95 - 0.95) | 0.89 (0.89 - 0.90) | 0.40 (0.35 - 0.44) | 0.87 (0.87 - 0.88) | 0.60 (0.59 - 0.62) | 0.90 (0.89 - 0.91) | **0.73 (0.42 - 1.03)** |
| **Pathway** | 0.66 (0.59 - 0.74) | 0.82 (0.78 - 0.86) | 0.82 (0.78 - 0.85) | 0.53 (0.47 - 0.58) | -0.62 (-.76 - -.48) | 0.52 (0.43 - 0.61) | -0.26 (-.47 - -.05) | 0.28 (0.15 - 0.41) | **0.07 (-0.82 - 0.68)** |
| **ME+GE +sex** | 0.56 (0.43 - 0.69) | 0.81 (0.77 - 0.85) | 0.83 (0.77 - 0.89) | 0.53 (0.41 - 0.64) | -0.42 (-.63 - -.20) | 0.52 (0.38 - 0.65) | -0.2 (-0.47 - 0.07) | 0.26 (0.11 - 0.41) | **0.09 (-0.54 - 0.72))** |
| **ME+GE+ sex per gene** | 0.01 (-1.11 - 1.13) | 0.84 (0.71 - 0.97) | 0.67 (0.47 - 0.87) | 0.26 (- 0.98 - 1.5) | -4.22 (-10.57 - 2.14) | 0.75 (0.67 - 0.84) | -0.48 (-1.33 - 0.37) | 0.58 (0.09 - 1.07) | **0.37 (-0.8 - 1.54)** |

| | | | | | | | | | |
|---|---|---|---|---|---|---|---|---|---|
| **Deeper (3 densely connected)** | 0.81 (0.79 - 0.82) | 0.84 (0.84 - 0.85) | 0.87 (0.86 - 0.87) | 0.76 (0.74 - 0.77) | -0.18 (-0.32 - -.05) | 0.56 (0.54 - 0.59) | -0.27 (-0.33 - -.21) | 0.58 (0.56 - 0.61) | **0.14 (-0.63 - 0.91)** |

*Supplementary Table 5. Results for the base network and variations to the base network. Age prediction performance in explained variance ($R^2$) for each cohort in a cohort-wise cross validation, mean over 10 runs with 95% confidence interval.*

| Gene | Mean percentage fold 1 | Mean percentage fold 2 | Mean percentage fold 3 | Mean percentage fold 4 | Mean all folds |
|---|---|---|---|---|---|
| COL11A2 | 0.57 | 0.55 | 0.34 | 0.34 | 0.61 |
| AFAP1 | 0.85 | 0.59 | 0.53 | 0.53 | 0.60 |
| OTUD7A | 0.37 | 0.32 | 0.63 | 0.63 | 0.55 |
| PTPRN2 | 0.28 | 0.80 | 0.43 | 0.43 | 0.51 |
| ADARB2 | 0.49 | 0.59 | 0.40 | 0.40 | 0.48 |
| CD34 | 0.20 | 0.42 | 0.64 | 0.64 | 0.45 |
| AGAP1 | 0.43 | 0.44 | 0.52 | 0.52 | 0.44 |
| IRS2 | 0.47 | 0.24 | 0.42 | 0.42 | 0.43 |
| DPYSL4 | 0.37 | 0.19 | 0.75 | 0.75 | 0.42 |
| CACNA1I | 0.59 | 0.37 | 0.35 | 0.35 | 0.41 |
| DNAJB6 | 0.59 | 0.39 | 0.21 | 0.21 | 0.40 |
| FBXO31 | 0.46 | 0.44 | 0.17 | 0.17 | 0.38 |
| GRM2 | 0.33 | 0.36 | 0.70 | 0.70 | 0.35 |
| PPP2R2D | 0.00 | 0.38 | 0.39 | 0.39 | 0.34 |
| MGMT | 0.08 | 0.33 | 0.31 | 0.31 | 0.29 |
| CHMP6 | 0.14 | 0.31 | 0.32 | 0.32 | 0.29 |

*Supplementary Table 6. Genes with contributions higher than 1% of the total weight for biological age prediction for three out of the four folds.*

#### V. LDL-level prediction

| LDL-level prediction | | | | | | | | | |
|---|---|---|---|---|---|---|---|---|---|
| | $R^2$ validation cohorts (excluding named cohort) | | | | $R^2$ (test cohort) | | | | Mean over all cohorts |
| | RS | LL | LLS | NTR | RS | LL | LLS | NTR | |
| **ME+GE** | 0.14 (0.13 - 0.15) | 0.13 (0.12 - 0.14) | 0.17 (0.16 - 0.18) | 0.05 (0.05 - 0.05) | 0.01 (-.00 - 0.01) | 0.07 (0.05 - 0.08) | -0.01 (-0.02 - 0.0) | 0.01 (0.01 - 0.02) | **0.02 (-0.04 - 0.08)** |
| **ME** | -0.4 (-0.61 - -0.18) | -0.3 (-0.45 - -0.14) | -0.32 (-.56 - -0.08) | -0.38 (-.54 - -0.23) | -0.56 (-.85 - -.28) | -0.42 (-.65 - -.19) | -0.31 (-.55 - -.06) | -0.38 (-.54 - -.21) | **-0.56 (-1.38 - 0.26)** |
| **GE** | 0.13 (0.12 - 0.13) | 0.12 (0.12 - 0.13) | 0.16 (0.16 - 0.17) | 0.05 (0.05 - 0.06) | -0.01 (-.01 - -.00) | 0.04 (0.04 - 0.05) | -0.01 (-0.01 - -.00) | -.06 (-.08 - - .04) | **-0.02 (-0.09 - 0.06)** |
| **ME+GE (L1 ME)** | 0.12 (0.11 - 0.13) | 0.11 (0.09 - 0.12) | 0.14 (0.12 - 0.15) | 0.05 (0.04 - 0.07) | 0.00 (-0.02 - 0.03) | 0.05 (0.04 - 0.06) | -0.05 (-.07 - -.02) | 0.02 (-.00 - .03) | **-0.00 (-0.07 - 0.06)** |
| **ME+GE (L1 GE)** | 0.10 (0.09 - 0.12) | 0.08 (0.06 - 0.10) | 0.09 (0.07 - 0.10) | 0.04 (0.03 - 0.05) | -0.03 (-0.06 - -.01) | 0.02 (0.01 - 0.03) | -0.01 (-0.02 - 0.01) | -0.03 (-.05 - -.01) | **-0.01 (-0.04 - 0.03)** |
| **Deeper (3 densely connected)** | 0.11 (0.07 - 0.15) | 0.01 (-.01 - 0.02) | 0.12 (0.10 - 0.14) | 0.04 (0.03 - 0.05) | 0.0 (-.01 - .01) | -0.0 (-0.01 - 0.00) | -0.0 (-.01 - 0.01) | 0.01 (0.00 - 0.01) | **-0.00 (-0.01 - 0.01)** |
| **Deeper (Pathway)** | 0.11 (0.10 - 0.11) | 0.11 (0.10 - 0.11) | 0.11 (0.10 - 0.11) | 0.05 (0.05 - 0.06) | -0.03 (-.04 - -.010) | 0.03 (0.02 - 0.03) | -0.0 (-.01 - 0.00) | -0.01 (-.02 - 0.00) | **-0.01 (-0.04 - 0.02)** |
| **ME+GE +sex** | -0.49 (-.93 - -0.06) | 0.02 (-0.03 - 0.08) | -1.34 (-1.66 - -1.01) | -2.22 (-2.76 - -1.68) | -0.79 (-1.68 - 0.1) | -0.03 (-.10 - 0.04) | -1.59 (-1.98 - -1.20) | -2.22 (-2.81 - -1.62) | **-1.60 (-3.35 - 0.15)** |

*Supplementary Table 7. Performance for cohort-wise cross validation for LDL level prediction, mean with 95% CI over 10 runs. Explained variance of 0 is equal to just predicting the mean. Explained variance can be negative when the network overfits and, consequently, has poorer predictive performance than predicting the mean in the test set.*

#### Supplementary Figures

##### VI. Deeper networks

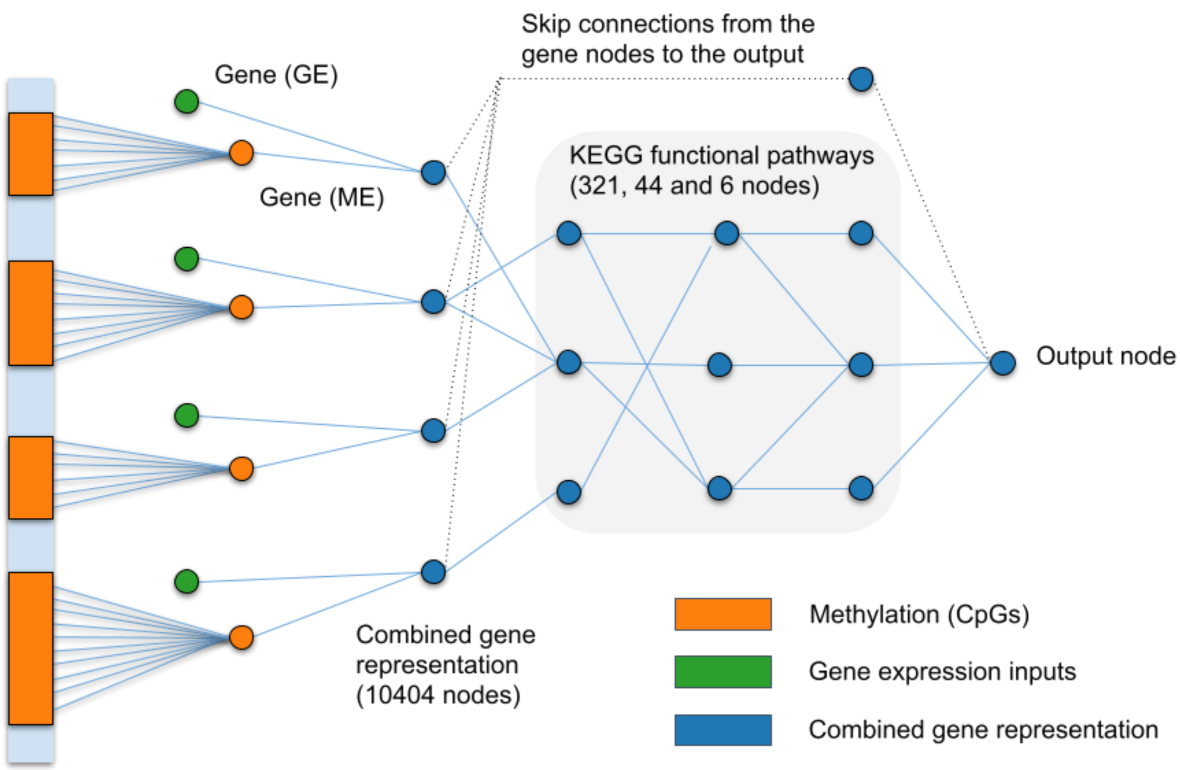


*Supplementary Figure 1. Overview of the pathway neural network. Using the ME+GE as a basis the genes are grouped and connected to their corresponding pathways. KEGG's functional pathways have 3-layer hierarchical structure with 321, 44 and 6 nodes each. The densely connected neural has the same number of neurons as the pathway network. In the densely connected network, the KEGG layers are replaced by fully connected layers where each node is connected to all nodes in previous and subsequent layers.*

#### VII. Smoking: pathway importance

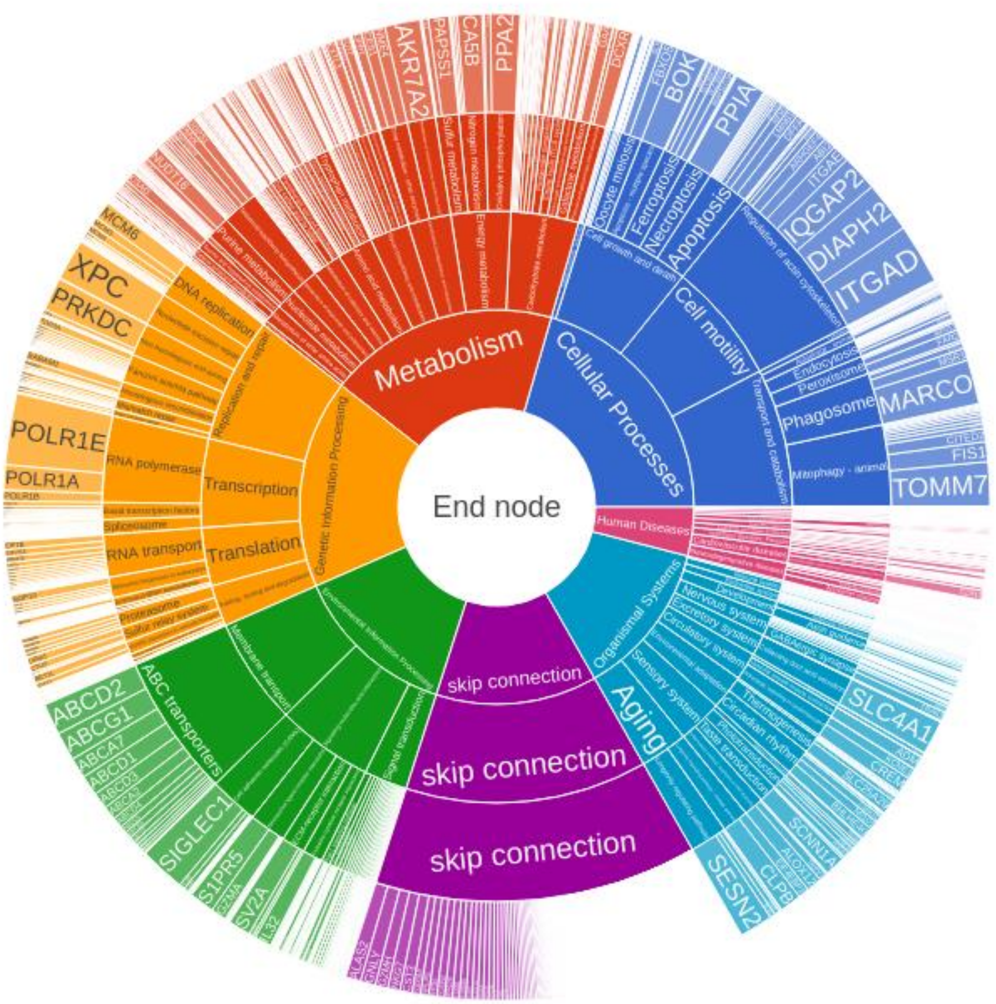


*Supplementary Figure 2. Importance of KEGG functional pathways and their corresponding genes for predicting smoking status. Skip connections connect each gene right away to the end node to ensure that each gene is connected to the output.*

### VIII. Smoking status: omic-specific information

Simulated data for subtype analysis

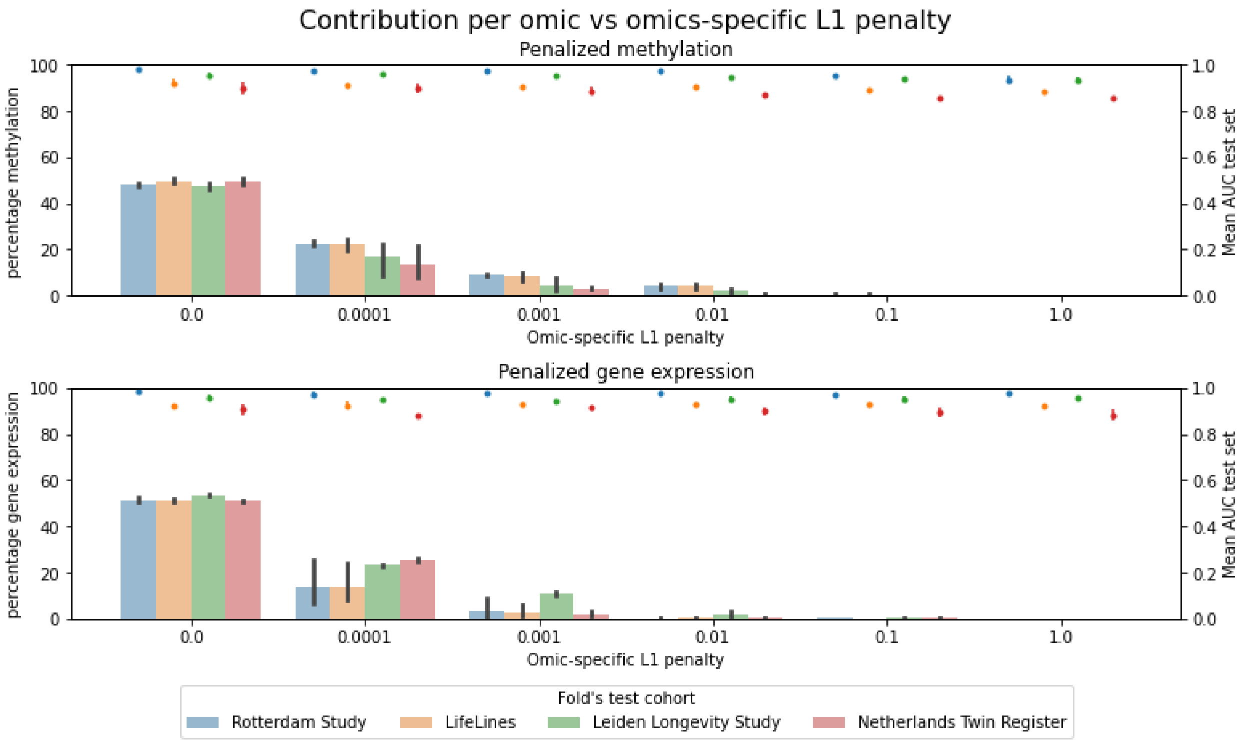


*Supplementary Figure 3. Contribution per omic versus omics-specific L1 penalty. The bar plots show the percentage of the total weight of the penalized omics. Additionally, the mean AUC and standard deviation is denoted by the colored points using the right axis.*

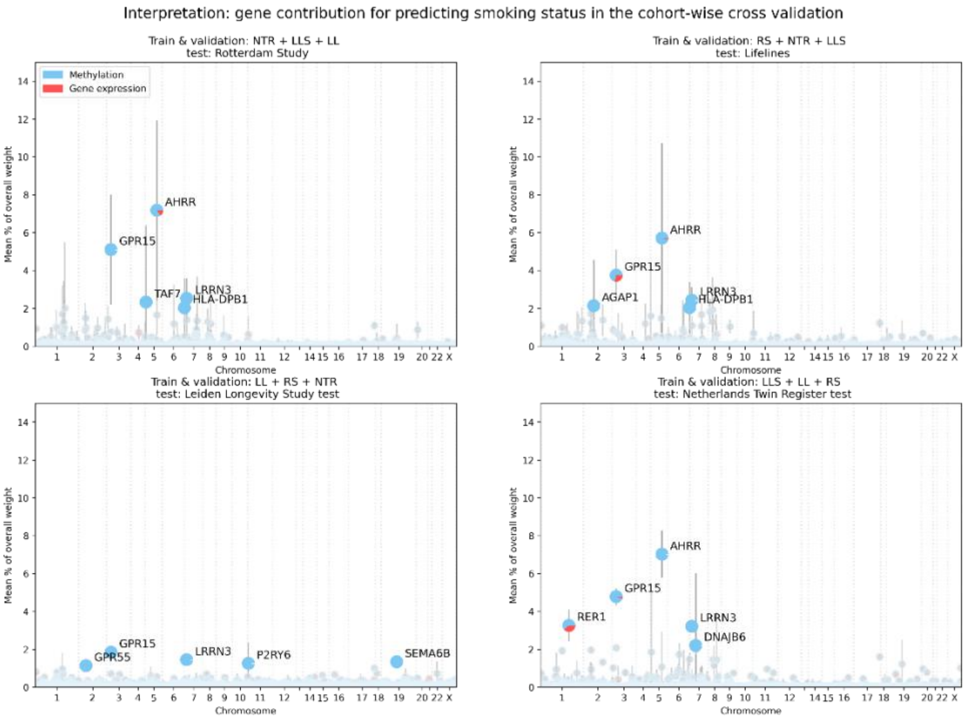


*Supplementary Figure 4. Smoking prediction with the ME+GE network penalized for gene expression genes with an omic-specific L1 penalty of 0.001.*

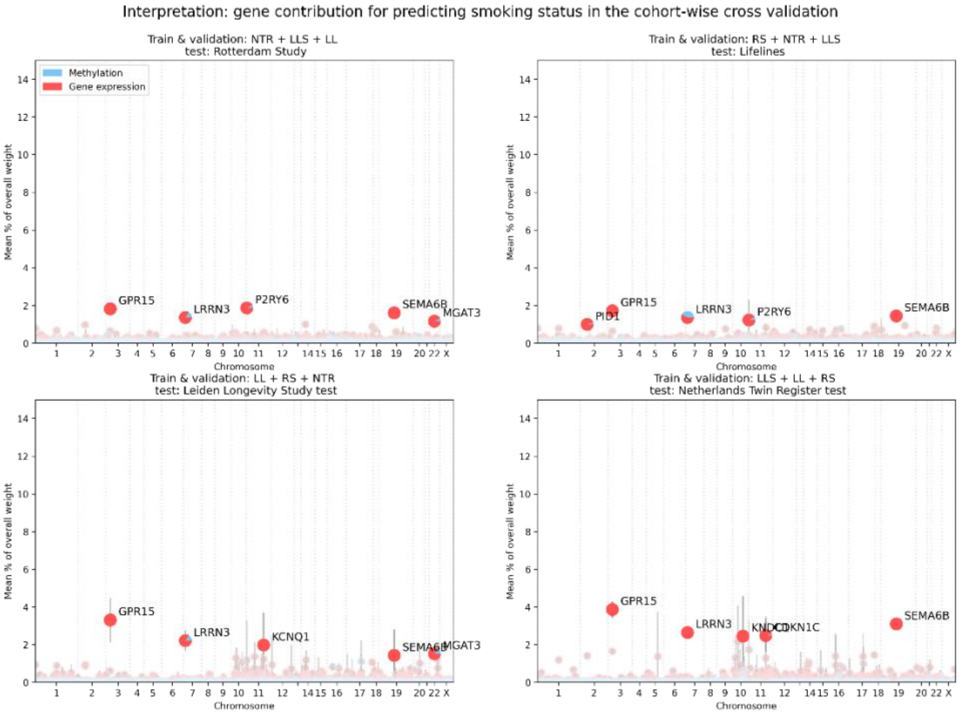


*Supplementary Figure 5. Smoking prediction with the ME+GE network penalized for methylation genes with an omic-specific L1 penalty of 0.001.*

### IX. Age prediction

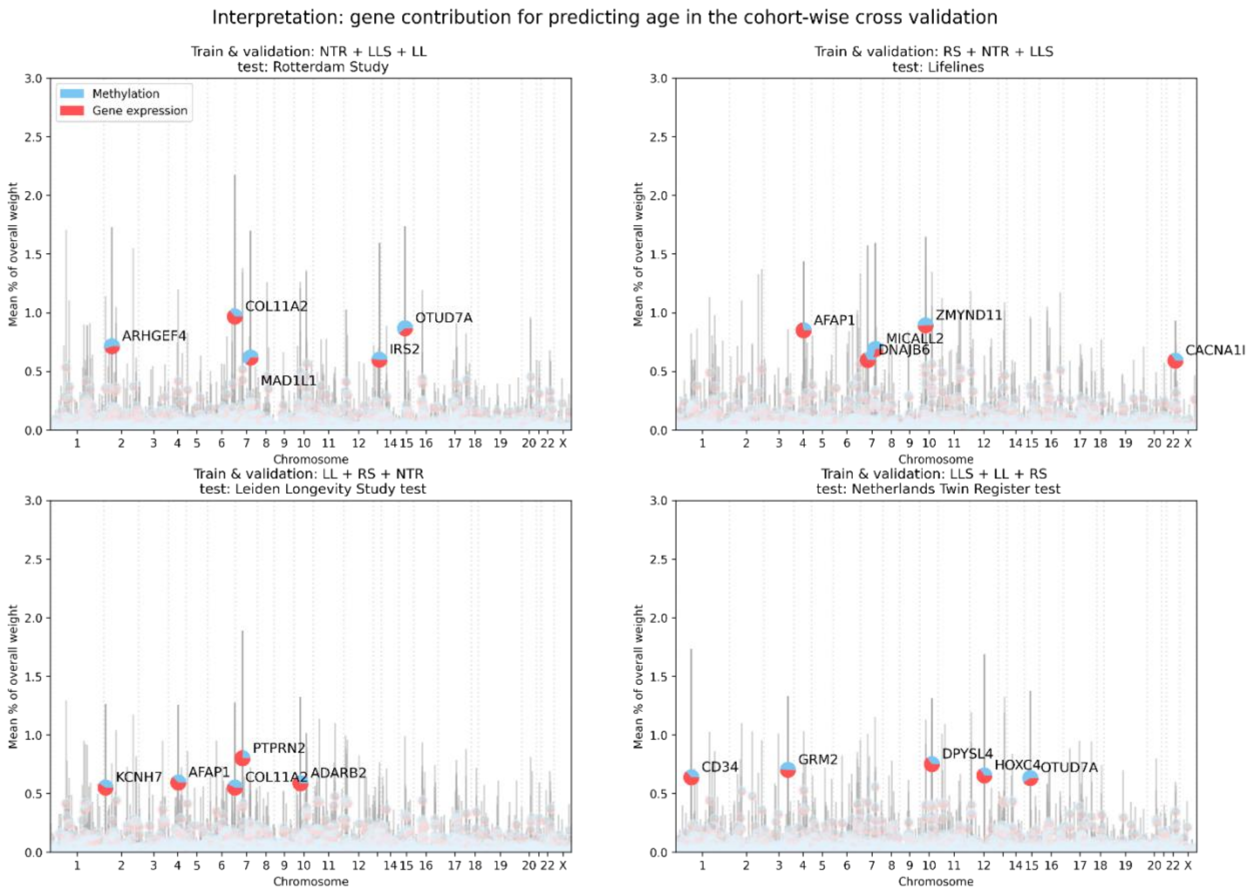


*Supplementary Figure 6. Interpratation for networks trained to predict biological age. Mean and standerd deviation over ten runs with different random seed for each fold. Percent of the weights occupied per gene, higher prercentages are more important for predicting biological age.*

### X. Age: sex specific effects

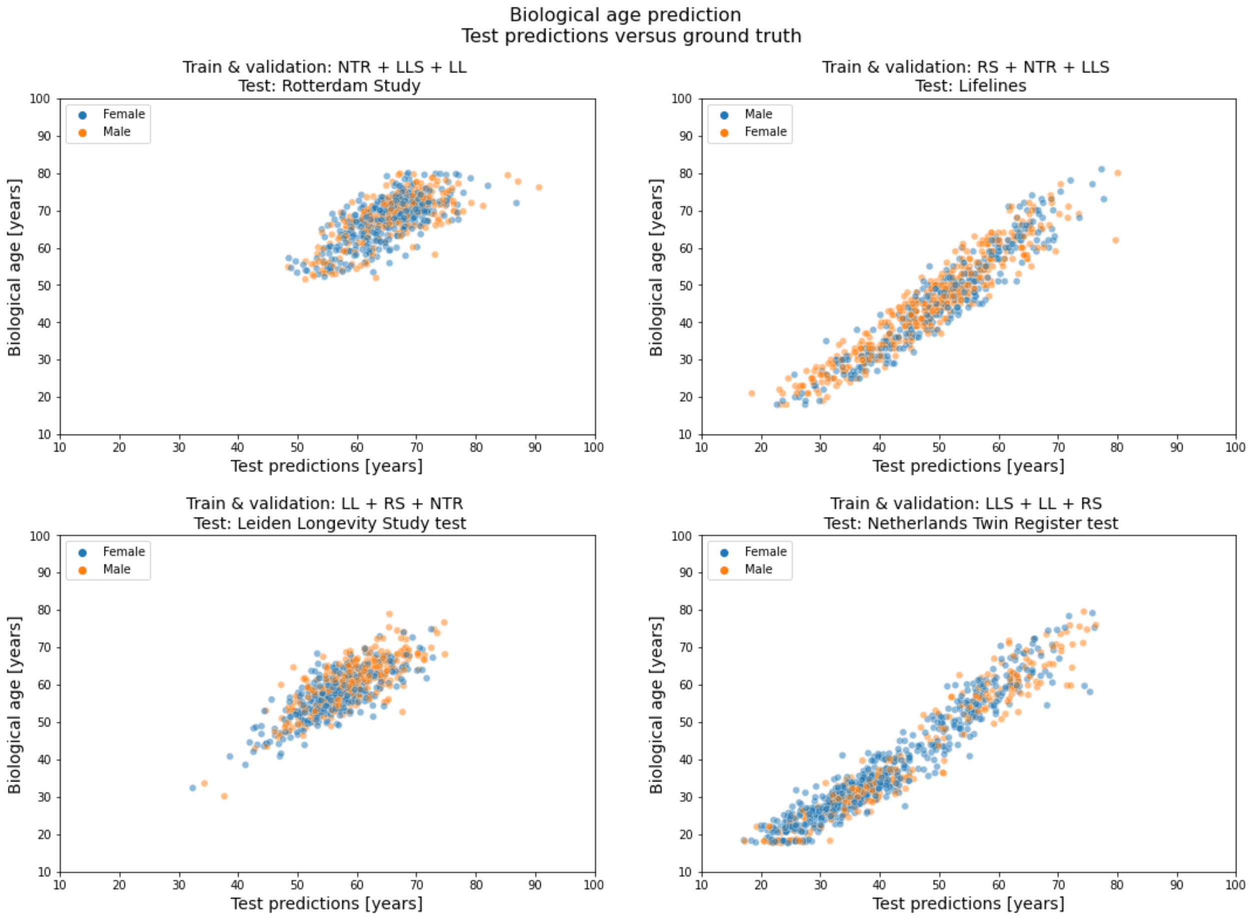


*Supplementary Figure 7: Biological age prediction versus the ground truth for each of the test cohorts.*

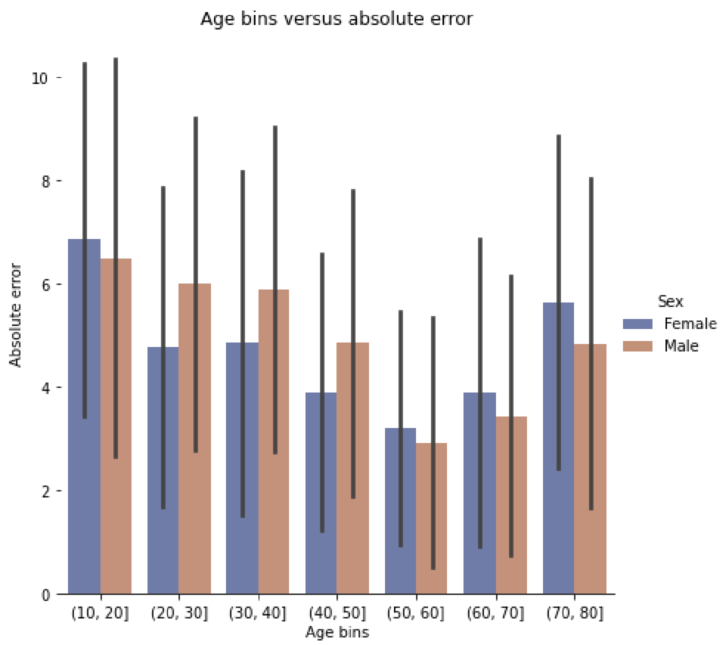


*Supplementary Figure 8. Absolute error for biological age prediction binned with a 10-year interval.*

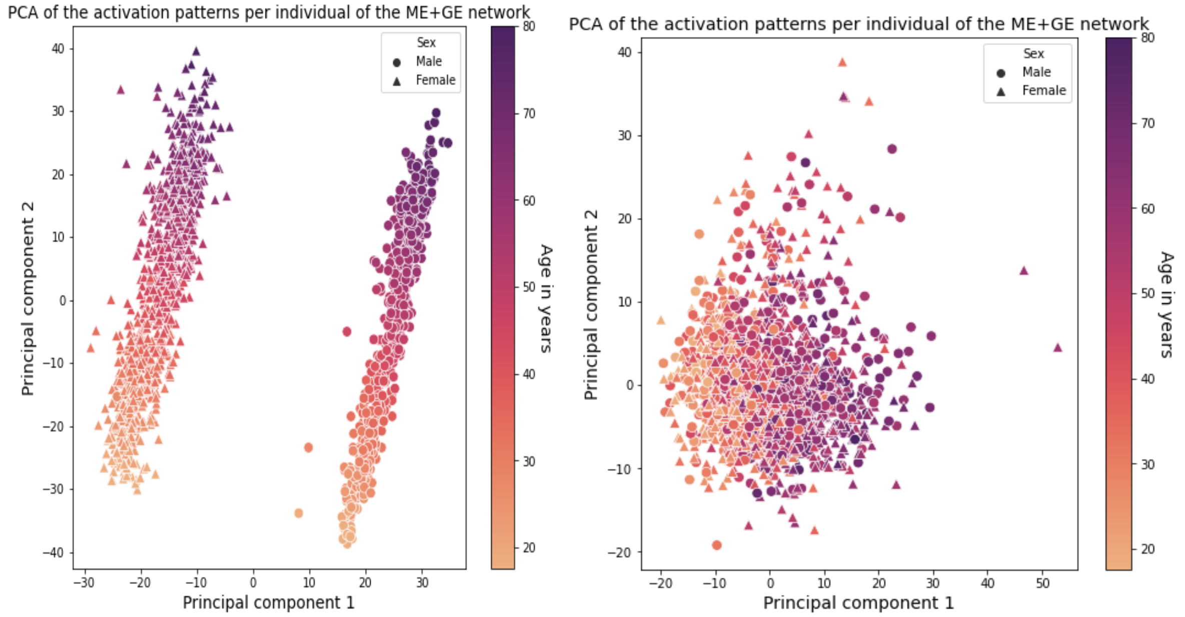


*Supplementary Figure 9. PCA of the activation excluding all genes on the X chromosome. Left including genes on the X-chromosome, right excluding genes on the X-chromosome.*

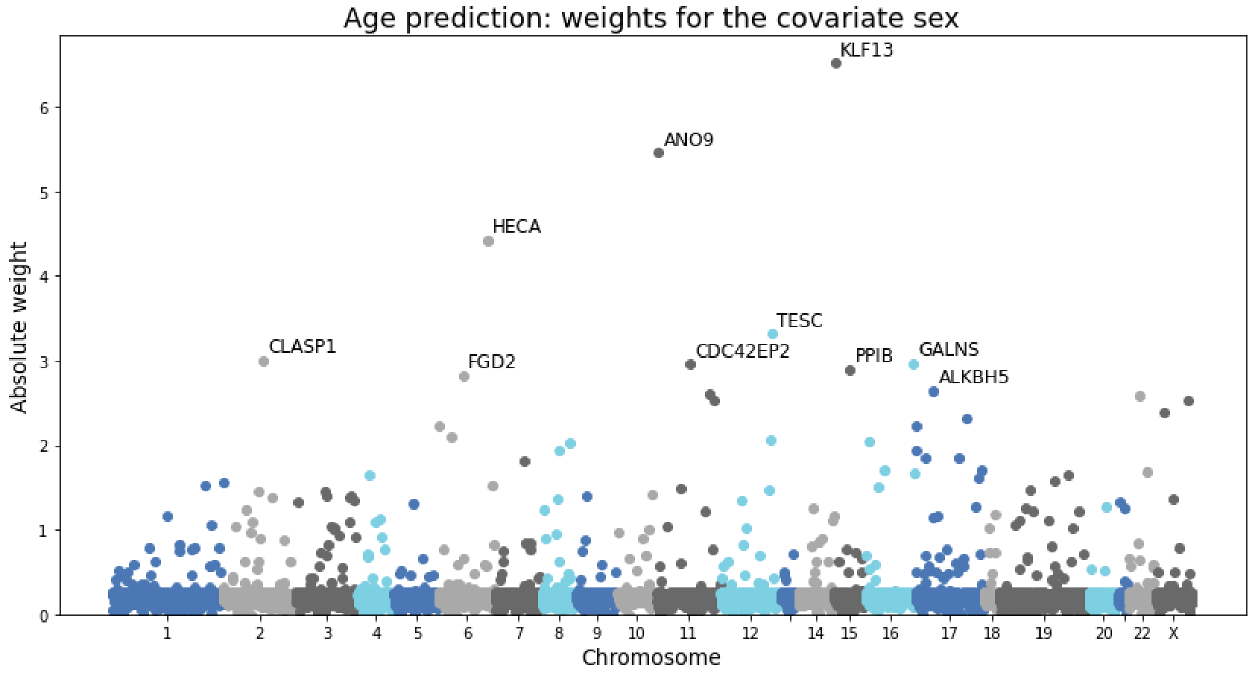


*Supplementary Figure 10. Weights for the covariate sex for each gene. Higher weights suggest that there is a larger difference between the sexes.*

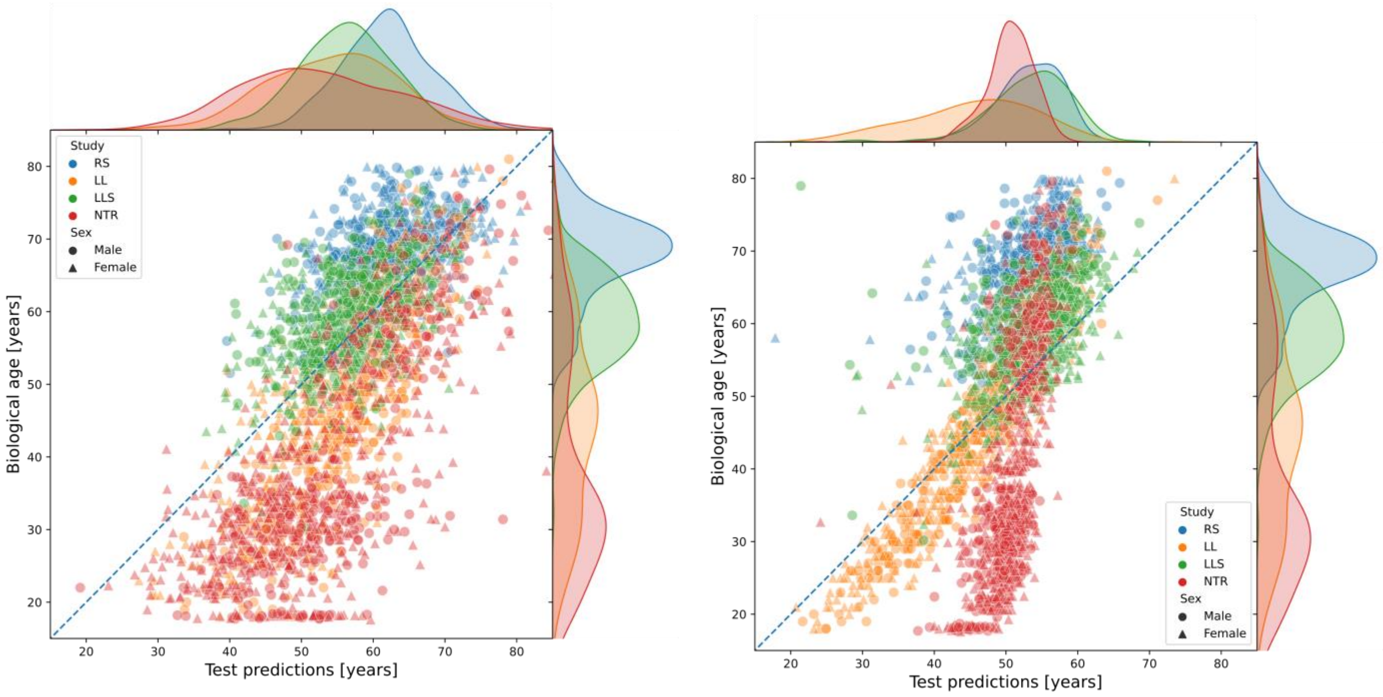


*Supplementary Figure 11 and 12. Test predictions for each cohort with their corresponding distributions for the GE and ME network, respectively.*

#### XI. Age: omic specific effects

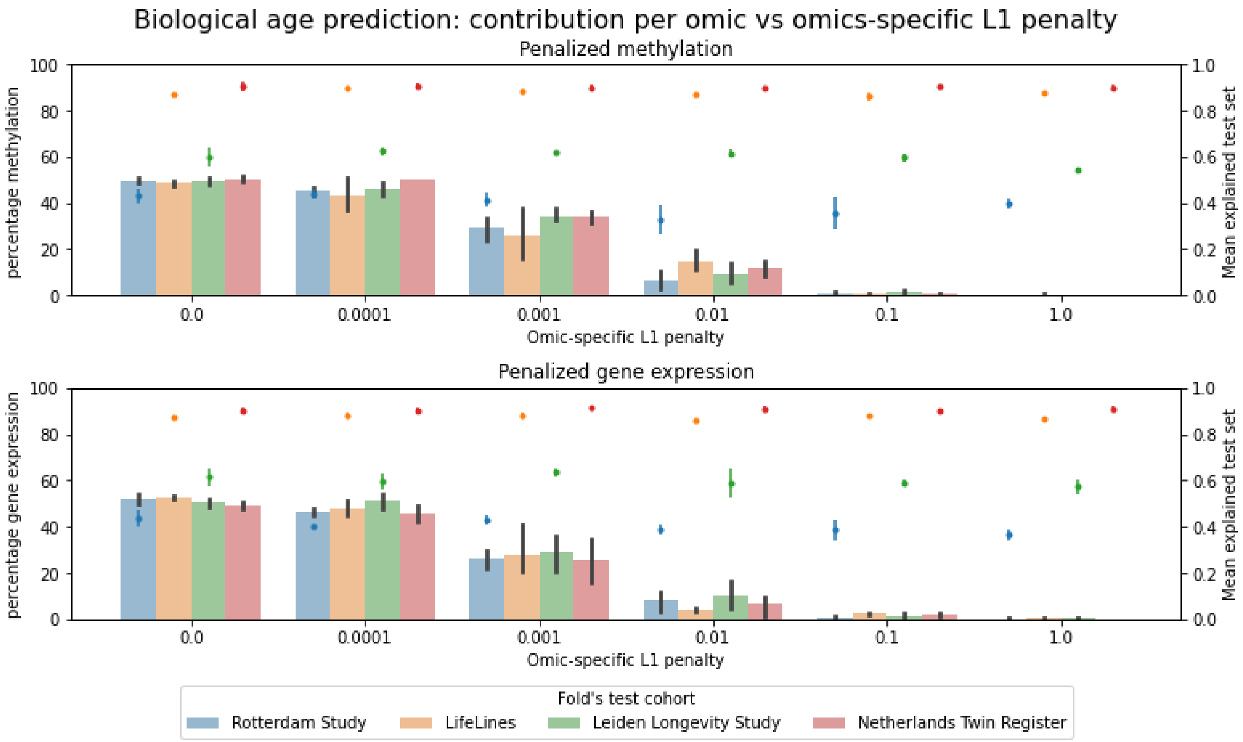


*Supplementary Figure 13. Omic specific penalty and its effect on predictive performance and the total percentage of the weight associated with that omic.*

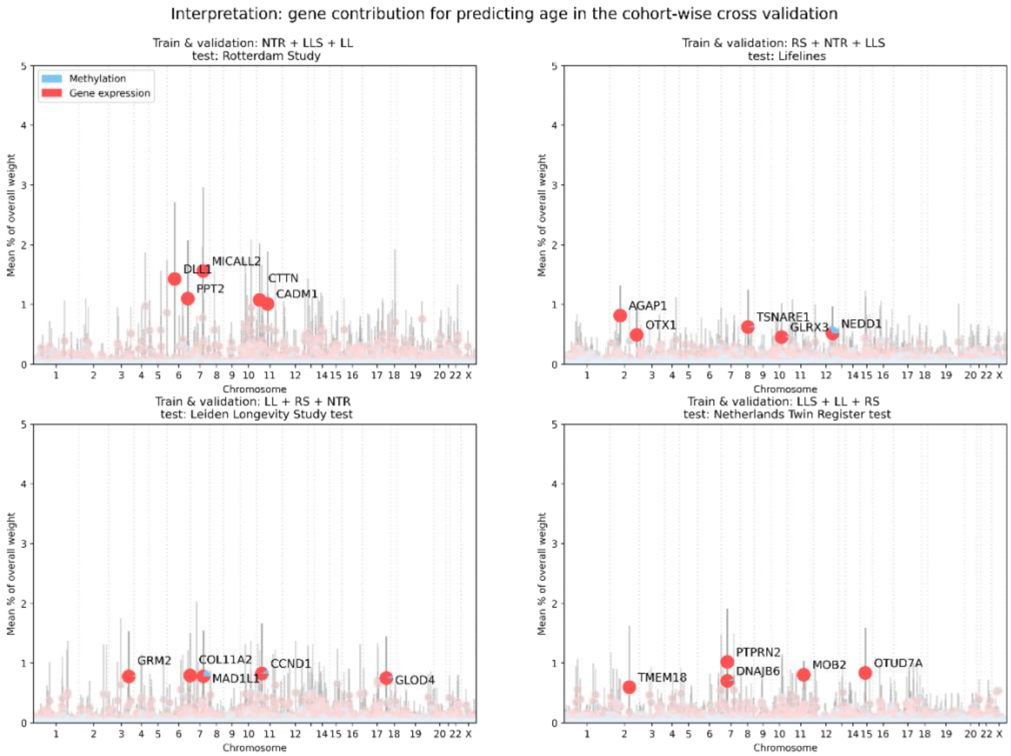


*Supplementary Figure 14. Biological age prediction with an omic-specific L1 penalty of 0.01 for the methylation input.*

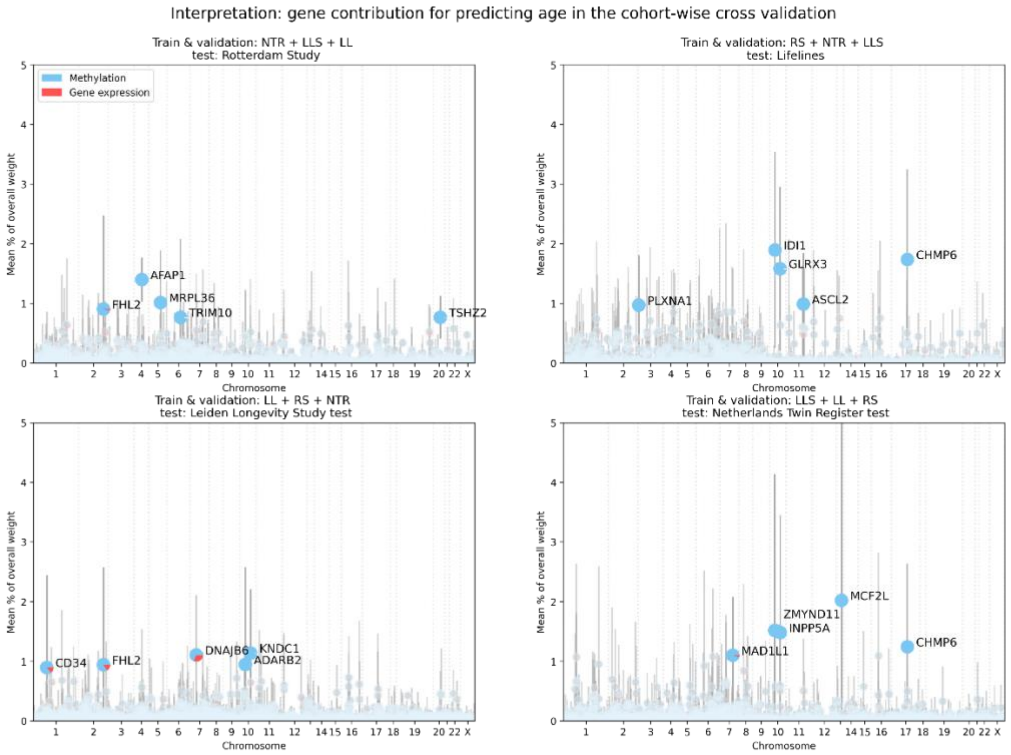


*Supplementary Figure 15. Biological age prediction with an omic-specific L1 threshold of 0.01 for gene expression input.*

#### XII. Age: pathway importance

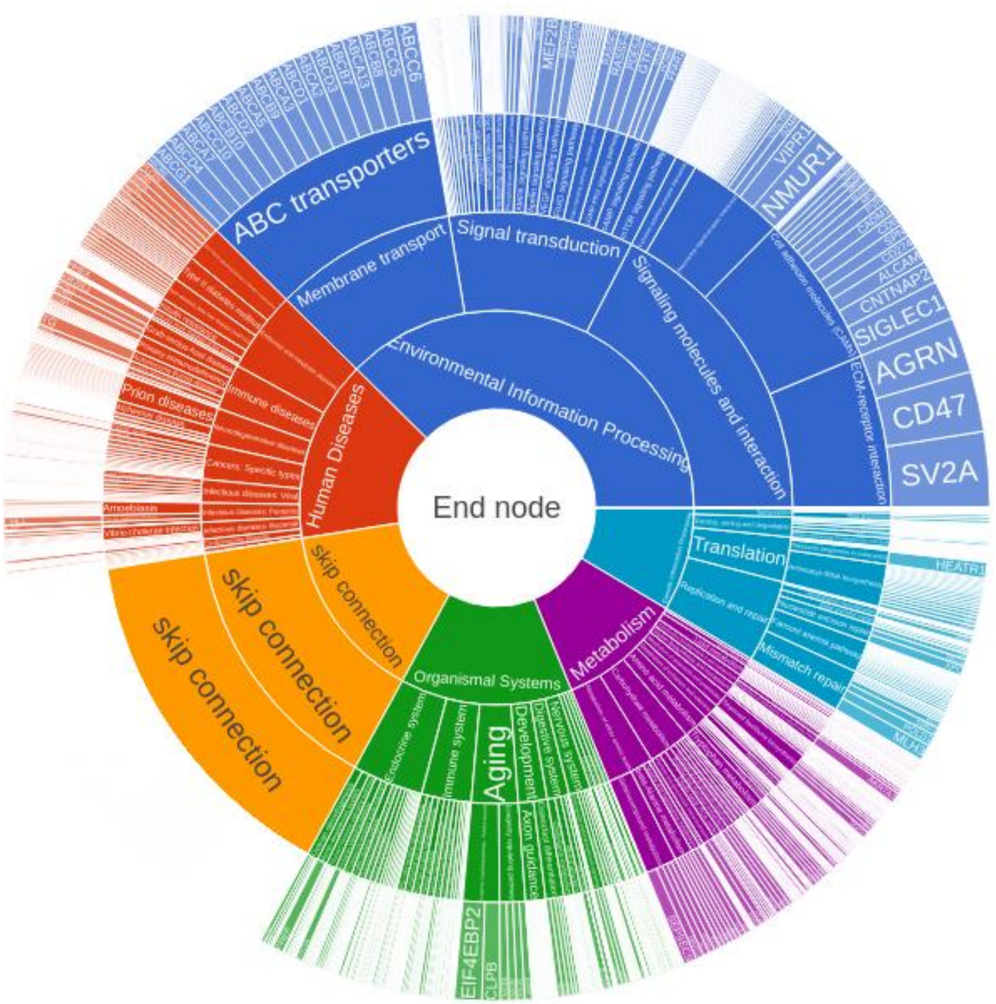


Supplementary Figure 16. *Importance of KEGG functional pathways and their corresponding genes for biological age. Skip connections connect each gene right away to the end node to ensure that each gene is connected to the output*

#### XIII. LDL prediction

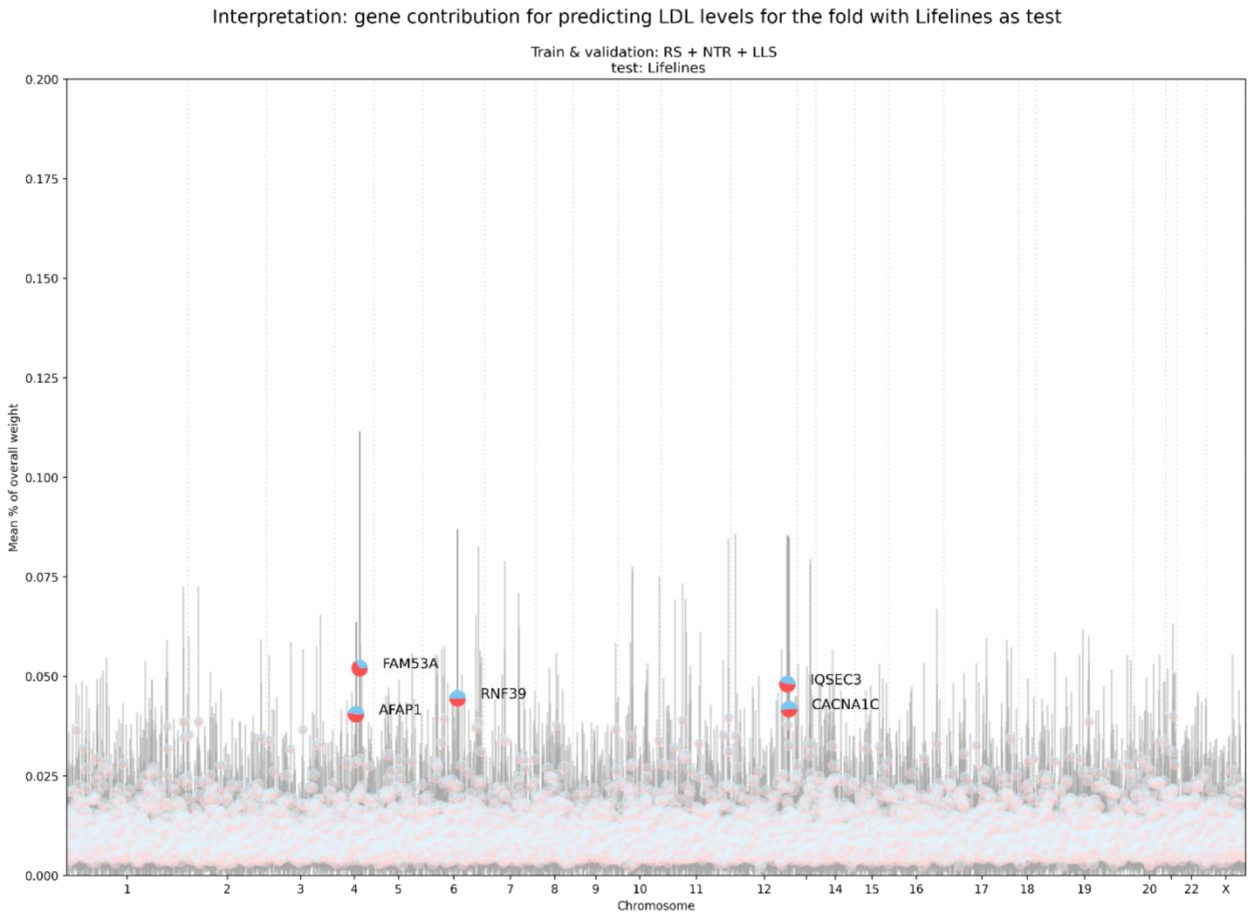


*Supplementary Figure 17. Weights for each gene as a percentage of the total weight in the network to predict LDL levels. Only the second fold was predictive and is shown here. Note that the small scale of the y-axis, all weights are close to zero.*
